# Reduced dopaminergic reinforcement, not learning capacity, limits operant learning in aging *Drosophila*

**DOI:** 10.64898/2026.06.17.732983

**Authors:** Ishrat Jahan, Helen Holvoet, Jean-François De Backer, Ilona C. Grunwald Kadow

## Abstract

Aging is associated with a progressive decline in cognitive function, including the ability to adapt behavior based on its consequences. While classical conditioning in *Drosophila melanogaster* has provided key insights into reinforcement learning, how aging impacts operant learning and adjusting behavior based on action outcomes remains unclear. Here, we used a closed-loop optogenetic paradigm to test how aging affects operant learning and the role of dopaminergic neurons (DANs; PPL1 and PAM) in this process. Activation of distinct DAN subsets revealed that both young and aged flies retain PPL1-dependent avoidance learning, indicating preserved action–outcome learning with age. However, learning in aged flies depended on prolonged reinforcement: unlike young flies, they failed to learn under shorter optogenetic stimulation durations, suggesting an aging-associated reduced dopaminergic reinforcement rather than a loss of learning capacity. Moreover, reducing mitochondrial antioxidant capacity via SOD2 knockdown in PPL1 neurons phenocopied the aging-related deficit, implicating oxidative stress in impaired reinforcement signaling. In contrast, broad PAM neuron activation drove robust learning across ages and stimulation regimes. Nevertheless, functional dissection of PAM subpopulations revealed subtype-specific aging-associated vulnerability within dopaminergic circuits. In line with the behavioral data, PPL1 but not PAM neurons exhibited age-dependent reductions in cell size. Together, our findings suggest that aging selectively reduces dopaminergic reinforcement in a DAN subtype-dependent manner while preserving the capacity for operant learning. Increasing reinforcement length rescues this deficit, indicating that altered dopaminergic signaling, rather than impaired learning capacity, is a key driver of age-related cognitive decline.

## Introduction

Aging is associated with progressive decline in cognitive function, alongside broader changes in sensory processing and motor performance (López-Otín et al., 2013; Murman, 2015). However, cognitive aging is not uniform. Crystallized abilities, reflecting accumulated knowledge, remain relatively stable with age, whereas working memory, the capacity to temporarily maintain and manipulate information to adjust ongoing behavior, declines with age (Blazer et al., 2015; Salthouse, 1994). The mechanism why some forms of memory remain relatively stable with age while others decline remains incompletely understood. This question can be addressed in *Drosophila melanogaster*, a genetically tractable model that combines a short lifespan with precise access to defined neural populations, enabling direct investigation of underlying neural circuits (Griffith, 2012; Piper & Partridge, 2018). In *Drosophila*, working memory–like processes can be modeled using operant conditioning in which an animal modifies its behavior based on the outcome of its own actions, i.e., rewards reinforce actions, while punishments suppress them (Murphy & Lupfer, 2014; Rajagopalan et al., 2023; Wiggin et al., 2021).

Like in other animals, dopaminergic neurons (DANs) in flies convey reinforcement signals that shape behavior through modulation of activity within the mushroom body (MB), a higher-order associative center (Aso, Sitaraman, et al., 2014; Liu et al., 2012; Mao & Davis, 2009; Perisse et al., 2016; Waddell, 2013). Distinct DAN populations encode opposing outcomes: protocerebral posterior lateral 1 (PPL1) neurons are primarily associated with punishment, whereas protocerebral anterior medial (PAM) neurons are mostly linked to reward (Aso, Hattori, et al., 2014; Siju et al., 2020). Through this circuit, dopaminergic signaling links recent outcomes to subsequent behavioral responses, biasing behavior toward avoidance or attraction. With aging, neuronal circuits underpinning learning and memory become increasingly affected by oxidative stress and mitochondrial dysfunction, which compromise neuronal integrity and particularly impact dopaminergic signaling (Brooker et al., 2024; Trist et al., 2019; Watanabe et al., 2024). While the effects of aging on classical conditioning have been studied in *Drosophila melanogaster* (Tamura et al., 2003; Tonoki et al., 2020; Tonoki & Davis, 2012), and the central role of dopaminergic reinforcement in this form of learning is well established in young adults (Aso et al., 2010; Liu et al., 2012), much less is known about how aging influences operant conditioning and the underlying dopaminergic reinforcement processes. Given the precise tools and knowledge of the neuronal connectivity, the *Drosophila* model system could provide additional insights also for higher animals including humans.

To test whether working memory–like processes that depend on dopaminergic reinforcement can be captured under operant conditions and how they are affected by aging, we used the optoPAD system as an operant learning assay (Moreira et al., 2019). This closed-loop platform couples feeding behavior to temporally precise activation of defined DAN populations, such that reinforcement depends directly on the animal’s own actions. Behavior is quantified as changes in interaction with the stimulated food source, revealing avoidance or attraction. We examined learning under two different stimulation durations (prolonged and short) using optogenetically-induced reinforcement via different, genetically labeled, DAN subsets and additionally tested the contribution of oxidative stress by expressing the mitochondrial Superoxide dismutase 2 (SOD2) RNAi in PPL1 neurons. Our findings suggest that aging does not abolish operant learning but instead weakens dopaminergic reinforcement signaling. This deficit is selective for specific dopaminergic circuits, with PPL1 neurons showing greater vulnerability than PAM neurons. Evidence further indicates that cellular stress mechanisms contribute to these changes, pointing to altered dopaminergic signaling rather than a loss of learning capacity as the primary driver of age-related cognitive decline. Together, our data reveal how aging affects operant learning and working memory–like processes that depend on dopaminergic reinforcement.

## Results

### Dopaminergic PPL1 neurons drive operant avoidance behavior depending on stimulation duration and age

We first examined whether aging alters baseline feeding behavior to assess the suitability of the flyPAD feeding assay for the analysis of aging-related performance in operant learning. In the flyPAD assay (Itskov et al., 2014), the cumulative number of sips and total sip number were measured in young (7–8 days) and aged (28–30 days) wild-type Canton-S (WT-CS) flies presented with 5% sucrose. Feeding behavior was comparable between age groups, with no detectable difference in cumulative number of sips over time or total sip number at 60 min (Fig. 1A), indicating that baseline feeding capacity is preserved with age.

**Figure 1.**
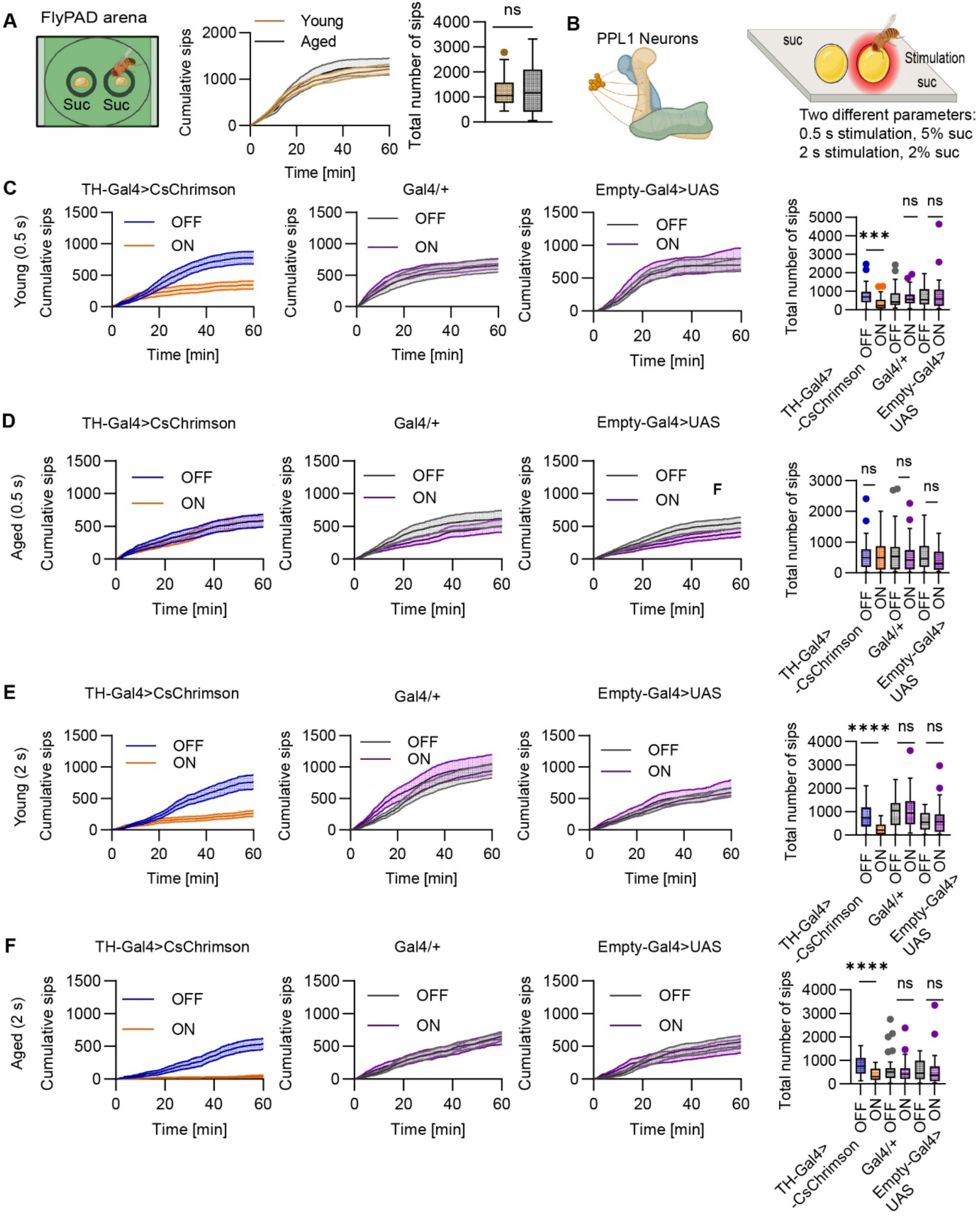
Dopaminergic PPL1 neurons drive operant avoidance behavior depending on stimulation duration and age. (A) Baseline sucrose feeding behavior measured using the flyPAD assay. Schematic of the flyPAD arena, in which individual flies are presented with a single 5% sucrose food source (left). Young (7–8 days) and aged (28–30 days) WT-CS flies were presented with 5% sucrose following 24 h starvation. Cumulative sip number over 60 min (middle) and total sip number at 60 min (right). No significant differences were observed between age groups (Mann-Whitney U Test, p value = 0.7654). Baseline (WT-CS): young n = 34, aged n = 28. (B) Schematic of the closed-loop optoPAD paradigm used to drive operant avoidance learning. PPL1 DANs project to the vertical lobes (α/α’) and γ lobes of the mushroom body. Flies expressing CsChrimson in PPL1 neurons were tested under two stimulation conditions: 0.5 s red-light pulses paired with 5% sucrose, or 2 s red-light pulses paired with 2% sucrose. (C–D) Effect of short optogenetic stimulation (0.5 s, 5% sucrose). Cumulative sip number over 60 min (left) and total sip number at 60 min (right) under light OFF and ON conditions in young (C) and aged (D) flies for TH-Gal4>CsChrimson experimental flies (young n = 31, aged n = 29), Gal4/+ controls (young n = 29, aged n = 30), and Empty-Gal4>UAS controls (young n = 30, aged n = 29). (E–F) Effect of longer optogenetic stimulation (2 s, 2% sucrose). Cumulative sip number over 60 min (left) and total sip number at 60 min (right) under light OFF and light ON conditions in young (E) and aged (F) flies for TH-Gal4>CsChrimson experimental flies (young n = 29, aged n = 29), Gal4/+ controls (young n = 28, aged n = 30), and Empty-Gal4>UAS controls (young n = 31, aged n = 29). Light OFF (blue; grey) and light ON (orange; purple). Statistical comparisons were performed using the Wilcoxon matched-pairs signed-rank test. (C) Young (0.5 s): TH-Gal4>CsChrimson, p = 0.0002; Gal4/+, p = 0.5903; Empty-Gal4>UAS, p = 0.9395. (D) Aged (0.5 s): TH-Gal4>CsChrimson, p = 0.2770; Gal4/+, p = 0.6963; Empty-Gal4>UAS, p = 0.0857. (E) Young (2 s): TH-Gal4>CsChrimson, p < 0.0001; Gal4/+, p > 0.9999; Empty-Gal4>UAS, p = 0.9154. (F) Aged (2 s): TH-Gal4>CsChrimson, p < 0.0001; Gal4/+, p = 0.4161; Empty-Gal4>UAS, p = 0.2560.

A decline in dopaminergic signaling has been implicated in aging-related cognitive decline (Bäckman et al., 2006) and in neurodegenerative diseases such as Parkinson’s disease (Brooker et al., 2024; Hirth, 2010; Watanabe et al., 2024). In *Drosophila*, previous studies have reported age-related reductions in dopaminergic function, including decreased dopamine levels and selective impairments in dopaminergic signaling (Neckameyer et al., 2000; Tonoki et al., 2020). However, recent work suggests that age-related changes in dopaminergic activity are more complex, as dopaminergic signaling can also become aberrantly elevated during memory consolidation in aged flies (Matsuno et al., 2026). Notably, studies examining the effects of aging on learning in *Drosophila* have focused primarily on classical (Pavlovian) conditioning paradigms, such as odor–sugar and odor–shock conditioning (Matsuno et al., 2026; Tonoki et al., 2020), leaving open how aging affects action-outcome related forms of learning, such as operant learning. To address whether operant learning declined during aging and to analyze the role of DANs in a possible decline, we established whether closed-loop optogenetic activation of DANs would drive operant learning when paired with a sugar reward. In the context of classical associative learning, PPL1 neurons, which are activated by electric shock or high temperature (Galili et al., 2014; Mao & Davis, 2009; Riemensperger et al., 2005), reinforce aversion of a previously neutral or positive sensory stimulus (Aso, Sitaraman, et al., 2014). To this end, CsChrimson, a red-shifted channelrhodopsin variant that depolarizes neurons upon red-light illumination (Klapoetke et al., 2014), was expressed in PPL1 neurons using the TH-Gal4 driver, which covers this DAN subset (Friggi-Grelin et al., 2003). The flies were given the choice between two drops of sugar (food source), with each interaction on the light-paired sucrose source triggering a red-light pulse, activating CsChrimson-expressing PPL1 neurons in real time (Fig. 1B; Moreira et al., 2019). Given that PPL1 neurons encode aversive reinforcement in classical conditioning, we predicted that their pairing with feeding in an operant paradigm would decrease engagement with the light-paired food source compared to the non-light paired food source.

As predicted, activation of PPL1 DANs for 0.5 s progressively reduced interaction with the light-paired food source in young flies, measured as fewer activity bouts on the flyPAD (Fig. S1A), with a corresponding reduction in cumulative and total sip number under light ON compared to light OFF conditions (Fig.1C). This indicates that young flies learned to avoid the food source paired with TH-Gal4 neuron activation. In contrast, aged flies did not show a detectable difference between light ON and light OFF conditions under the same stimulation parameters suggesting a failure to learn (Fig. 1D). Control genotypes were unaffected by light delivery in both age groups, confirming that the behavioral effect depends on optogenetic activation of these DANs.

To test whether the absence of learned avoidance in aged flies originates from reduced dopaminergic reinforcement or a loss of learning capacity in the downstream circuits, i.e. Kenyon cells (KCs) and mushroom body output neurons (MBONs), we increased the strength of the reinforcement signal by extending the DAN stimulation duration from 0.5 s to 2 s. This combination allowed us to test whether longer dopaminergic reinforcement could restore avoidance learning in aged flies. Under these conditions, activation of PPL1 neurons in young flies again reduced interaction with the light-paired food source (Fig. 1E). Critically, aged flies now appeared to learn and displayed a significant reduction in cumulative number of sips and total sip number under light ON conditions compared with light OFF (Fig. 1F), indicating that aged MB circuits remain capable of supporting avoidance learning when the dopaminergic reinforcement signal is sufficiently strong.

We further characterized the behavioral change by analyzing two feeding parameters (Fig. S1). Under short stimulation of 0.5 s, young flies showed a reduction in activity bout number, indicating fewer returns to the light-paired food source, alongside a significant change in feeding burst duration (Fig. S1A). In contrast, aged flies showed no detectable changes in either parameter (Fig. S1B). Under longer stimulation, activity bout number was reduced in both young and aged flies, consistent with decreased engagement with the food source. Feeding burst duration showed a significant reduction in young flies but remained unchanged in aged flies (Fig. S1C, D). These results suggest that dopaminergic activation primarily reduced engagement with the light-paired food source, and that this effect is restored in aged flies under longer stimulation. Overall, the microstructure data are consistent with the pattern observed in total sip counts as aged flies fail to show reduced engagement under brief stimulation but recover this response when dopaminergic reinforcement is strengthened.

Together, these results demonstrate that both young and aged flies are capable of operant avoidance learning driven by PPL1 DANs. However, aged flies require longer optogenetic reinforcement through DANs to drive this behavior, indicating an age-related reduction in efficacy of dopaminergic reinforcement rather than a loss of operant learning capacity.

### Reduction of the enzyme SOD2 in DANs abolishes operant avoidance learning

Having established that TH>CsChrimson-driven operant avoidance can be rescued by reinforcement strength in aged flies, we next asked whether this learning depends on the mitochondrial redox state of those DANs. Aging is broadly associated with mitochondrial dysfunction and elevated reactive oxygen species (ROS) (López-Otín et al., 2013), and DANs may be especially affected, as dopamine metabolism itself generates additional oxidative burden (Burbulla et al., 2017; Graham, 1978; Meiser et al., 2013) and natural variation in antioxidant capacity has been linked to age-related dopamine neuron decline in flies (Coleman et al., 2024). Consistent with this, mutations in the *Drosophila* orthologues of the Parkinson’s disease genes parkin and PINK1, which regulate mitochondrial quality control, lead to mitochondrial dysfunction and selective dopaminergic neuron degeneration in flies (Clark et al., 2006; Greene et al., 2003; Park et al., 2006). Moreover, we have previously shown that aging-dependent decline of SOD2, a mitochondrial antioxidant enzyme that limits oxidative stress by converting superoxide radicals into less reactive oxygen species (ROS), contributes strongly to the functional decline of olfactory projection neurons (Hussain et al., 2018). We therefore reasoned that mitochondrial antioxidant capacity within DANs may be required to support effective dopaminergic negative reinforcement signaling. To analyze operant learning through DANs with reduced SOD2 levels, we generated flies expressing not only UAS-CsChrimson but also SOD2 RNAi to knock down the enzyme via RNA interference (RNAi) selectively in TH-Gal4-positive neurons. These flies, young and old, were tested in the optoPAD assay using a 5% sucrose solution, in which interaction with the food source triggered 0.5 s red light stimulation (Fig. 2A).

**Figure 2.**
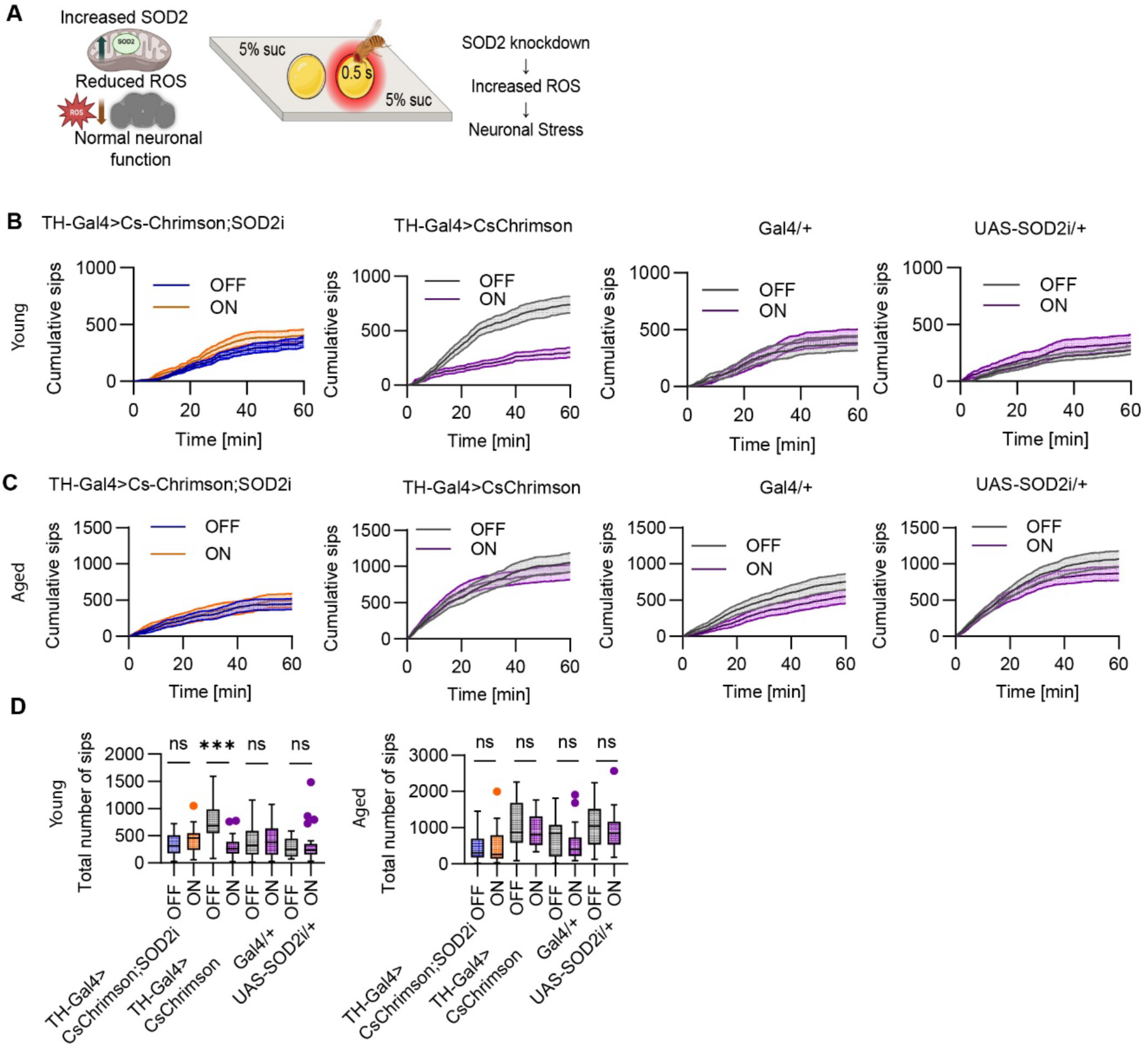
SOD2 knockdown in DANs abolishes operant avoidance behavior. (A) Schematic illustrating the role of SOD2 in regulating mitochondrial oxidative stress and the experimental paradigm. Reduction of SOD2 expression increases ROS levels and induces neuronal stress. Flies were tested in the optoPAD assay under closed-loop conditions, in which sip events on a 5% sucrose solution triggers 0.5 s red light activation of CsChrimson in TH-Gal4-positive DANs. (B–C) Effect of SOD2 knockdown on cumulative sip number over 60 min under light OFF and light ON conditions in young (B) and aged (C) flies. Data are shown for TH-Gal4>CsChrimson;SOD2i flies (young n = 21, aged n = 25), TH-Gal4>CsChrimson experimental flies (young n = 21, aged n = 22), Gal4/+ controls (young n = 21, aged n = 24) and UAS-SOD2i/+ controls (young n = 22, aged n = 28). (D) Total sip number at 60 min under light OFF and light ON conditions for the indicated genotypes in young (left) and aged (right) flies. Light OFF (blue; grey) and light ON (orange; purple). Statistical comparisons were performed using the Wilcoxon matched-pairs signed-rank test. (D) Young: TH-Gal4>CsChrimson;SOD2i, p= 0.3488; TH-Gal4>CsChrimson, p = 0.0002; Gal4/+, p = 0.6091; UAS-SOD2i/+, p = 0.4060. Aged: TH-Gal4>CsChrimson;SOD2i, p = 0.6011; TH-Gal4>CsChrimson, p = 0.5028; Gal4/+, p = 0.1355; UAS-SOD2i/+, p = 0.0571.

As seen before (Fig. 1C, D), activation of PPL1 DANs (TH>CsChrimson) in young flies again resulted in operant avoidance behavior. In comparison, young SOD2 RNAi expressing flies did not show reduced interaction with the light-paired food source suggesting that learning did not take place. Cumulative number of sips and total sip number were comparable between light ON and light OFF conditions, indicating loss of this avoidance response (Fig. 2B–D). Control genotypes showed no differences between light conditions. Aged SOD2 RNAi-expressing flies also showed no detectable learning just like the aged control flies (Fig. 2C, D) consistent with the hypothesis that SOD2 levels were already reduced during natural aging in older animals (Davie et al., 2018; Hussain et al., 2018). Feeding microstructure analysis revealed no light-dependent changes in activity bout number or feeding burst duration in SOD2 RNAi flies in either age group (Fig. S1E, F). One control group (UAS-SOD2i/+ in aged flies under 0.5 s stimulation) showed a significant reduction in activity bout number under light-OFF conditions (Fig. S1F); however, this effect was not observed in other control genotypes and therefore is unlikely to reflect a driver-dependent process.

Together, these findings demonstrate that mitochondrial SOD2 in TH-Gal4-positive DANs is required for PPL1-driven operant avoidance learning in young flies, highlighting the importance of mitochondrial antioxidant capacity for dopaminergic reinforcement signaling.

### PAM neuron reinforcement induces operant learning independent of age

While PPL1 DANs primarily signal aversive reinforcement (Aso et al., 2010; Aso, Sitaraman, et al., 2014; Masek et al., 2015; Siju et al., 2020), PAM DANs are classically associated with appetitive reinforcement and reward signaling in *Drosophila* (Burke et al., 2012; Liu et al., 2012; Yamagata et al., 2015). However, their role in operant learning, and whether it is altered by aging, remains unclear. To test this, we optogenetically activated PAM DANs using the driver R58E02-Gal4 that is specifically expressed in all PAM DANs (Liu et al., 2012) (Fig. 3A). Flies were tested under both stimulation conditions used previously (0.5 s and 2 s). Surprisingly, under short stimulation (0.5 s), activation of PAM neurons produced robust avoidance rather than appetitive learning of the light-paired food source, reflected by a reduced cumulative number of sips and total sip number under light ON compared with light OFF conditions in both young and aged flies (Fig. 3B, C). Control genotypes showed no light-dependent changes. To determine whether this effect persisted with longer activation, stimulation duration was extended to 2 s. PAM activation again reduced feeding interactions in both age groups (Fig. 3D, E), while control genotypes showed no light-dependent changes. Feeding parameters analysis revealed reduced activity bout duration and feeding burst duration in experimental flies under both stimulation conditions (Fig. S2A–D).

**Figure 3.**
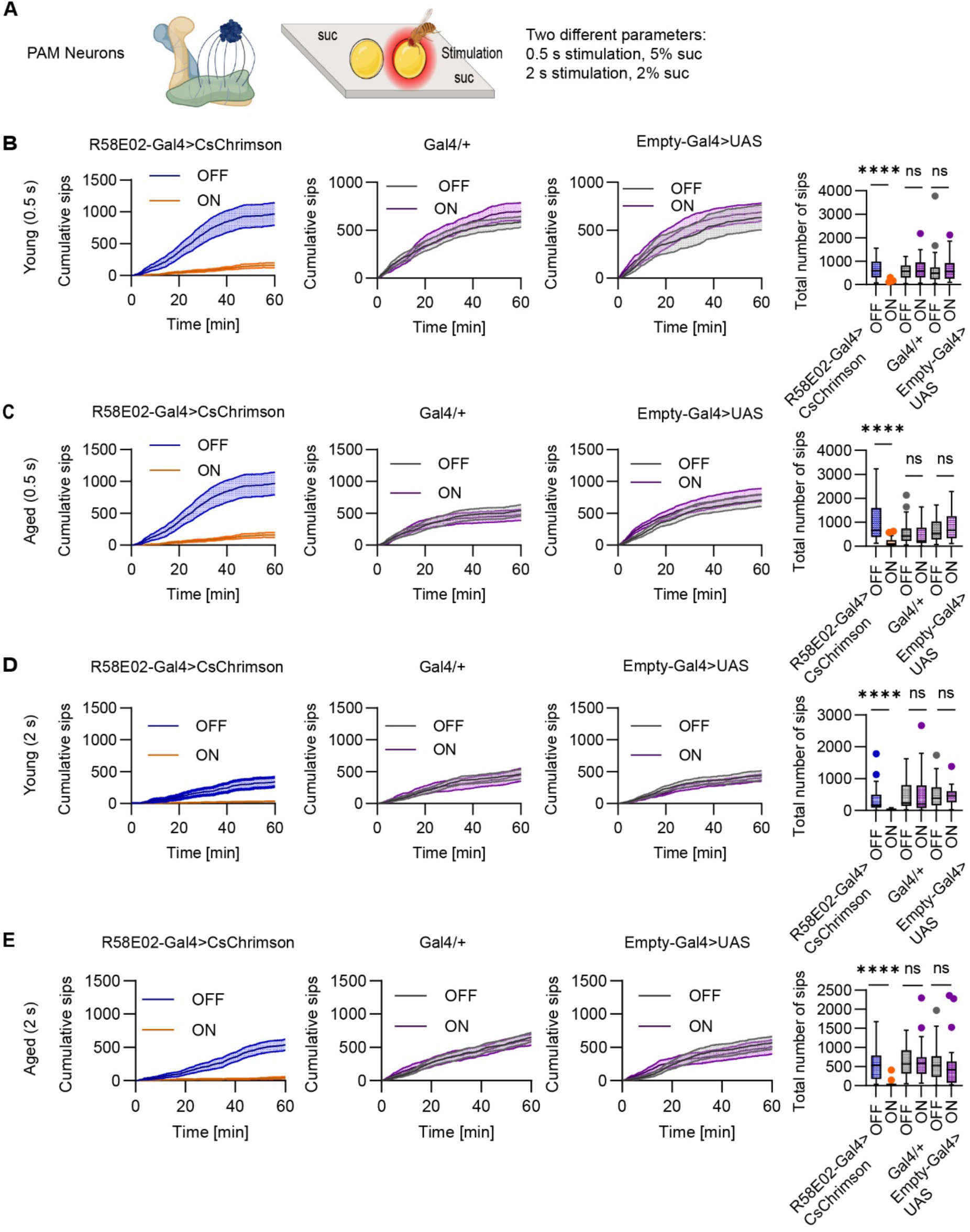
Activation of dopaminergic PAM neurons suppresses operant feeding behavior independently of stimulation duration and age. (A) Schematic of dopaminergic PAM neurons and optogenetic activation during the optoPAD assay under closed-loop conditions, in which sip events on the light-paired sucrose source triggered red light activation (0.5 s or 2 s). (B–C) Effect of short stimulation (0.5 s) on feeding behavior. Cumulative sip number over 60 min (left) and total sip number at 60 min (right) under light OFF and light ON conditions in young (B) and aged (C) flies. Data are shown for R58E02-Gal4>UAS-CsChrimson experimental flies (young n = 25, aged n = 22), R58E02-Gal4/+ controls (young n = 29, aged n = 30) and Empty-Gal4>UAS-CsChrimson controls (young n = 30, aged n = 28). (D–E) Effect of prolonged stimulation (2 s) on feeding behavior. Cumulative sip number over 60 min (left) and total sip number at 60 min (right) under light OFF and light ON conditions in young (D) and aged (E) flies. Data are shown for R58E02-Gal4>UAS-CsChrimson experimental flies (young n = 28, aged n = 21), R58E02-Gal4/+ controls (young n = 32, aged n = 30) and Empty-Gal4>UAS-CsChrimson controls (young n = 27, aged n = 27). Light OFF (blue; grey) and light ON (orange; purple). Statistical comparisons were performed using the Wilcoxon matched-pairs signed-rank test. (B) Young (0.5 s): R58E02>CsChrimson, p < 0.0001; Gal4/+, p = 0.7819; Empty-Gal4/UAS, p = 0.5186. (C) Aged (0.5 s): R58E02>CsChrimson, p < 0.0001; Gal4/+, p = 0.8512; Empty-Gal4/UAS, p = 0.5076. (D) Young (2 s): R58E02>CsChrimson, p < 0.0001; Gal4/+, p = 0.1600; Empty-Gal4/UAS, p = 0.8593. (E) Aged (2 s): R58E02>CsChrimson, p < 0.0001; Gal4/+, p = 0.6702; Empty-Gal4/UAS, p = 0.2609.

Together, these results show that broad PAM dopaminergic activation drives avoidance of the light-paired food source across stimulation durations. In contrast to PPL1-induced learning, PAM-reinforced operant learning was not affected by age.

### Subpopulation-specific effects of dopaminergic PAM-β′2a and β′2m/β′2p neurons on operant feeding behavior differ with age

Based on previous data, PAM neuron activation drives attractive behavior in naïve flies and leads to appetitive learning in classical odor-pairing experiments (Aso, Hattori, et al., 2014; Liu et al., 2012). Given that R58E02-Gal4 labels a heterogeneous PAM population that includes both appetitive and aversive subtypes and is larger in number as compared to PPL1 neurons (Yamagata et al., 2016; Musso et al., 2019; Lozada-Perdomo et al., 2025), we tested whether more restricted PAM subpopulations would drive operant approach behavior. We focused on the β’2 compartment, where DANs have been shown to mediate short-term appetitive reinforcement (Huetteroth et al., 2015; Lewis et al., 2015; Musso et al., 2019; Yamagata et al., 2016). We used two split-Gal4 lines: MB109B-Gal4, which labels PAM-β’2a neurons (Fig. 4A), and MB056B-Gal4, which labels neurons innervating the β’2m and β’2p subregions (Fig. 4D) (Aso, Hattori, et al., 2014). Flies expressing UAS-CsChrimson under these drivers were tested in the closed-loop optoPAD paradigm under both stimulation conditions used previously (0.5 s and 2 s). Given that β’2 PAM neurons encode appetitive reinforcement, we predicted that their activation would promote operant approach -increased engagement with the light-paired food source- rather than avoidance. Short stimulation (0.5 s) produced no detectable behavioral change in either driver line in young or aged flies (Fig. S3A, B and Fig. S4A, B). Therefore, we focused on prolonged 2 s stimulation, which did result in significant learning (Fig. 4).

**Figure 4.**
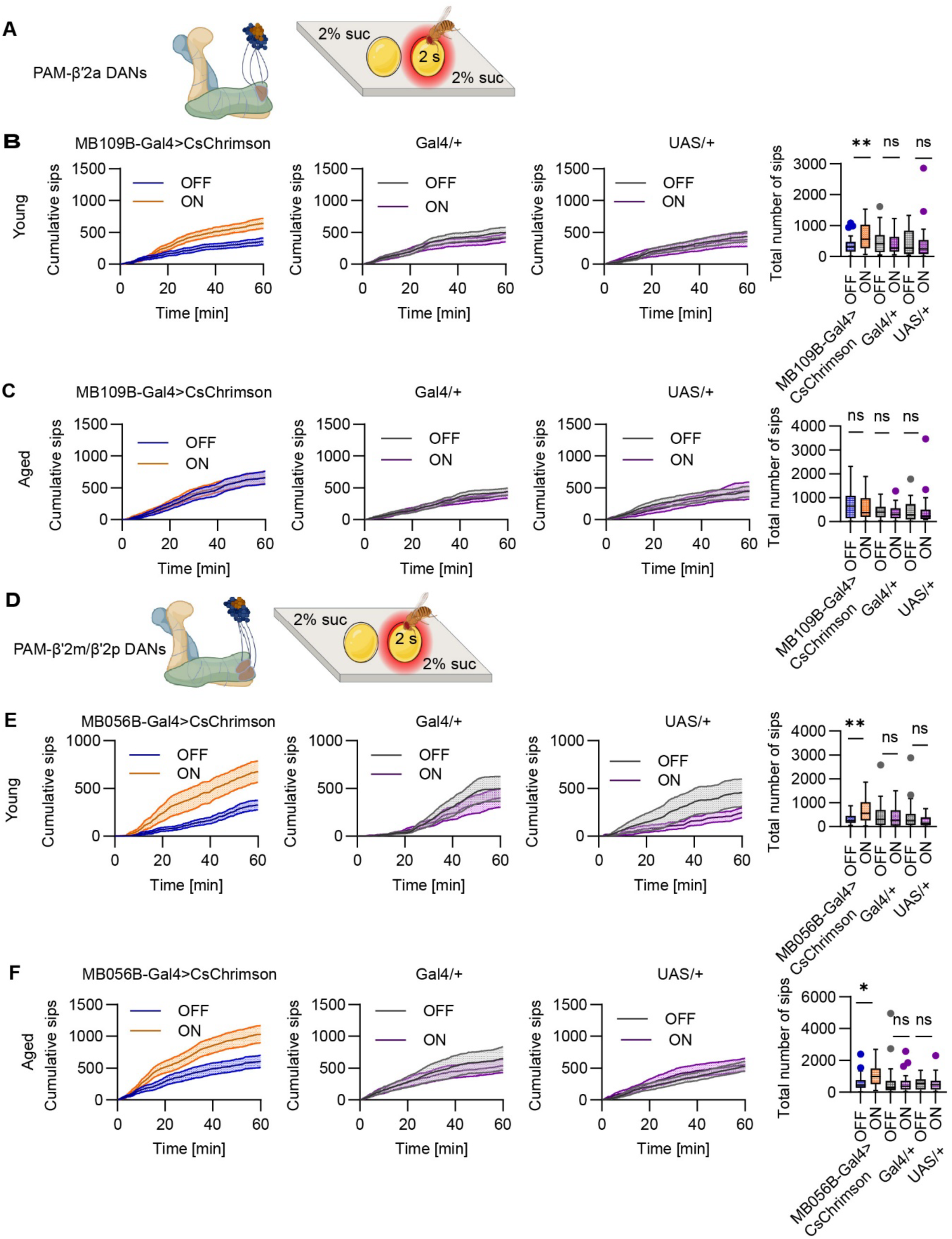
Subpopulation-specific effects of dopaminergic PAM-β′2a and β′2m/β′2p neurons on operant feeding behavior differ with age. (A, D) Schematic of dopaminergic PAM neuron subpopulations and the closed-loop optoPAD paradigm. (A) PAM-β′2a neurons (MB109B-Gal4) and (D) PAM-β’2m/β’2p neurons (MB056B-Gal4). Sip events on the light-paired sucrose source triggered 2 s red light stimulation (2.5 V), whereas interactions with the non-light-paired food source did not trigger stimulation. (B–C) Effect of prolonged optogenetic activation (2 s) of PAM-β′2a neurons.Cumulative sip number over 60 min (left) and total sip number at 60 min (right) under light OFF and light ON conditions in young (B) and aged (C) flies. Data are shown for MB109B-Gal4>CsChrimson experimental flies (young n = 29, aged n = 30), Gal4/+ controls (young n = 29, aged n = 32), and UAS/+ controls (young n = 27, aged n = 26). (E–F) Effect of prolonged optogenetic activation (2 s) of PAM-β’2m/β’2p neurons.Cumulative sip number over 60 min (left) and total sip number at 60 min (right) under light OFF and light ON conditions in young (E) and aged (F) flies. Data are shown for MB056B-Gal4>CsChrimson experimental flies (young n = 21, aged n = 28), Gal4/+ controls (young n = 21, aged n = 30), and UAS/+ controls (young n = 21, aged n = 26). Light OFF (blue; grey) and light ON (orange; purple). Statistical comparisons were performed using the Wilcoxon matched-pairs signed-rank test. (B) Young (2 s): MB109B-Gal4>CsChrimson, p = 0.0018; Gal4/+, p = 0.2867; UAS/+, p = 0.3212. (C) Aged (2 s): MB109B-Gal4>CsChrimson, p = 0.6702; Gal4/+, p = 0.3474; UAS/+, p = 0.7125. (E) Young (2 s): MB056B-Gal4>CsChrimson, p = 0.0049; Gal4/+, p = 0.8117; UAS/+, p = 0.2157. (F) Aged (2 s): MB056B-Gal4>CsChrimson, p = 0.0276; Gal4/+, p = 0.9233; UAS/+, p = 0.9552.

In contrast to reinforcement through the broad PAM lines, activation of PAM-β’2a neurons produced appetitive learning, i.e., increased feeding on the light-paired food source as compared to the non-light paired sucrose source (Fig. 4B). Interestingly, this learning was age-dependent (Fig. 4C). In young flies, MB109B-driven activation increased interaction with the light-paired food source, reflected by significantly higher cumulative number of sips and total sip number under light ON compared with light OFF conditions (Fig. 4B). This indicates that young flies learned to approach the food source paired with PAM-β’2a activation. In comparison, aged flies showed no light-dependent change and behaved like negative control genotypes that also showed no learning (Fig. 4C).

We next examined the slightly larger β’2m/β’2p population using MB056B-Gal4. In contrast to PAM-β’2a, MB056B activation increased feeding interactions in both young and aged flies (Fig. 4E, F), indicating that operant approach learning driven by this population is preserved with age. Control genotypes again showed no effect.

Feeding parameter analysis under prolonged stimulation revealed differential effects on feeding dynamics. In young flies, PAM-β’2a activation increased total activity bout duration without affecting feeding burst duration, whereas aged flies showed no detectable changes (Fig. S3E, F). In contrast, MB056B activation increased total activity bout duration in aged flies but not in young flies, while feeding burst duration remained unchanged (Fig. S4E, F). Under short stimulation, neither subpopulation produced detectable changes in feeding parameters (Fig. S3C, D and Fig. S4C, D). One control group (UAS-CsChrimson/+ in aged flies within the PAM-β’2a dataset) showed a significant decrease in total activity bout duration under light ON conditions (Fig. S3D); however, this effect was not present in other control genotypes and is therefore unlikely to reflect a driver-dependent process.

Together, these results show that distinct PAM subpopulations differentially support operant approach learning in an age-dependent manner. PAM-β’2a-driven approach is observed in young but not aged flies, whereas PAM-β’2m/β’2p -driven approach is preserved across both age groups, pointing to subpopulation-specific differences in how aging affects appetitive reinforcement signaling within the β’2 compartment.

### Age-related decline in soma size of PPL1 but not PAM neurons

Given the age- and cell type-specific differences in operant behavior observed in PAM and PPL1 neurons, we next examined whether these behavioral changes are accompanied by structural alterations in DANs. In previous work, we observed that aging as well as knock-down of SOD2 resulted in a decrease of soma size of olfactory projection neurons (Hussain et al., 2018). To assess cell number and cell size, GFP-labelled PPL1 and PAM neurons were visualized simultaneously using a combined TH,58E02-Gal4 driver (Siju et al., 2020) and imaged using confocal imaging in young and aged brains (Fig. 5A–B). We first quantified the number of GFP-expressing neurons in each population. No differences were observed between young and aged flies for either PPL1 or PAM neurons (Fig. 5C, D), indicating that dopaminergic neuron count is preserved with age and thus unlikely to be responsible for the observed decrease in PPL1-reinforced learning (White et al., 2010). We next measured the neuronal cell body surface by outlining individual somata and calculating their areas. PPL1 neurons showed a significant reduction in cell body size in aged compared to young flies (Fig. 5C), whereas PAM neurons showed no significant change with age (Fig. 5D). Across populations, PPL1 neurons exhibited larger cell body surface area than PAM neurons. Together, these results indicate that aging, to the here used stage, does not affect DAN number but manifests in a selective reduction in neuronal soma size of PPL1 neurons. This anatomical phenotype is in line with the aging-related learning defects described above.

**Figure 5.**
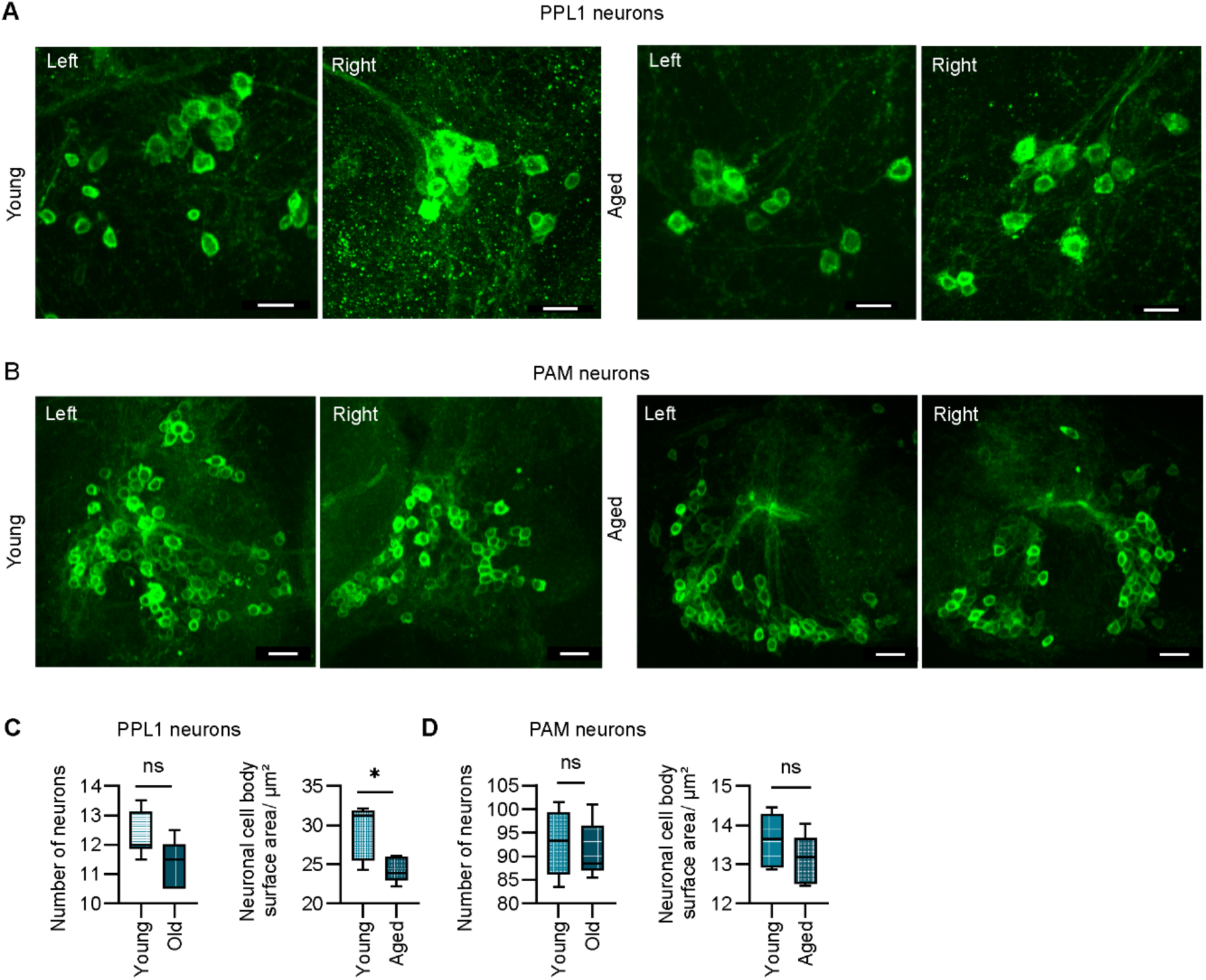
**Age-related changes in dopaminergic PPL1 and PAM neuron number and soma size in *Drosophila*** (A–B) Representative confocal images of GFP-expressing DANs in young and aged brains. Neurons were labelled using a combined TH,58E02-Gal4 driver, which labels both PPL1 (TH-Gal4) and PAM (R58E02-Gal4) clusters in the same brain. Representative images from left and right brain hemispheres are shown. Images are shown as total maximum intensity projections of confocal z-stacks. Scale bar = 10 µm. (C-D) Quantification of dopaminergic neuron number and cell body surface area (µm²). (C) PPL1 neurons: number of neurons (left) and neuronal cell body surface area (µm²; right) in young (n = 6) and aged (n = 5) flies. (D) PAM neurons: number of neurons (left) and neuronal cell body surface area (µm²; right) in young (n = 6) and aged (n = 5) flies. Cell body surface area (µm²) was measured by outlining individual neuronal somata. Statistical comparisons were performed using the Mann–Whitney U test. (C) PPL1: number of neurons, p = 0.2500; neuronal cell body surface area (µm²), p = 0.0303. (D) PAM: number of neurons, p = 0.7554; neuronal cell body surface area (µm²), p = 0.0625.

## Discussion

Cognitive abilities decline with aging. Whether this decline reflects an actual loss of learning capacity or a diminished sensitivity to contextual reinforcement remains difficult to disentangle. Here, we show that aged flies retain the capacity for operant learning, but they require longer dopaminergic reinforcement as compared to young flies. This suggests that aging reduces the strength of dopaminergic reinforcement signals rather than the ability to learn. Moreover, using optogenetic activation of distinct dopaminergic subsets as reinforcing signals as well as morphological analysis, we show that aging does not uniformly affect all DANs equally. These data are consistent with the observed differential loss of different types of aging-associated cognitive capacity and suggest that enhanced dopaminergic reinforcement, for instance, through more salient contextual signals, could improve cognitive performance (Fig. 6).

**Figure 6.**
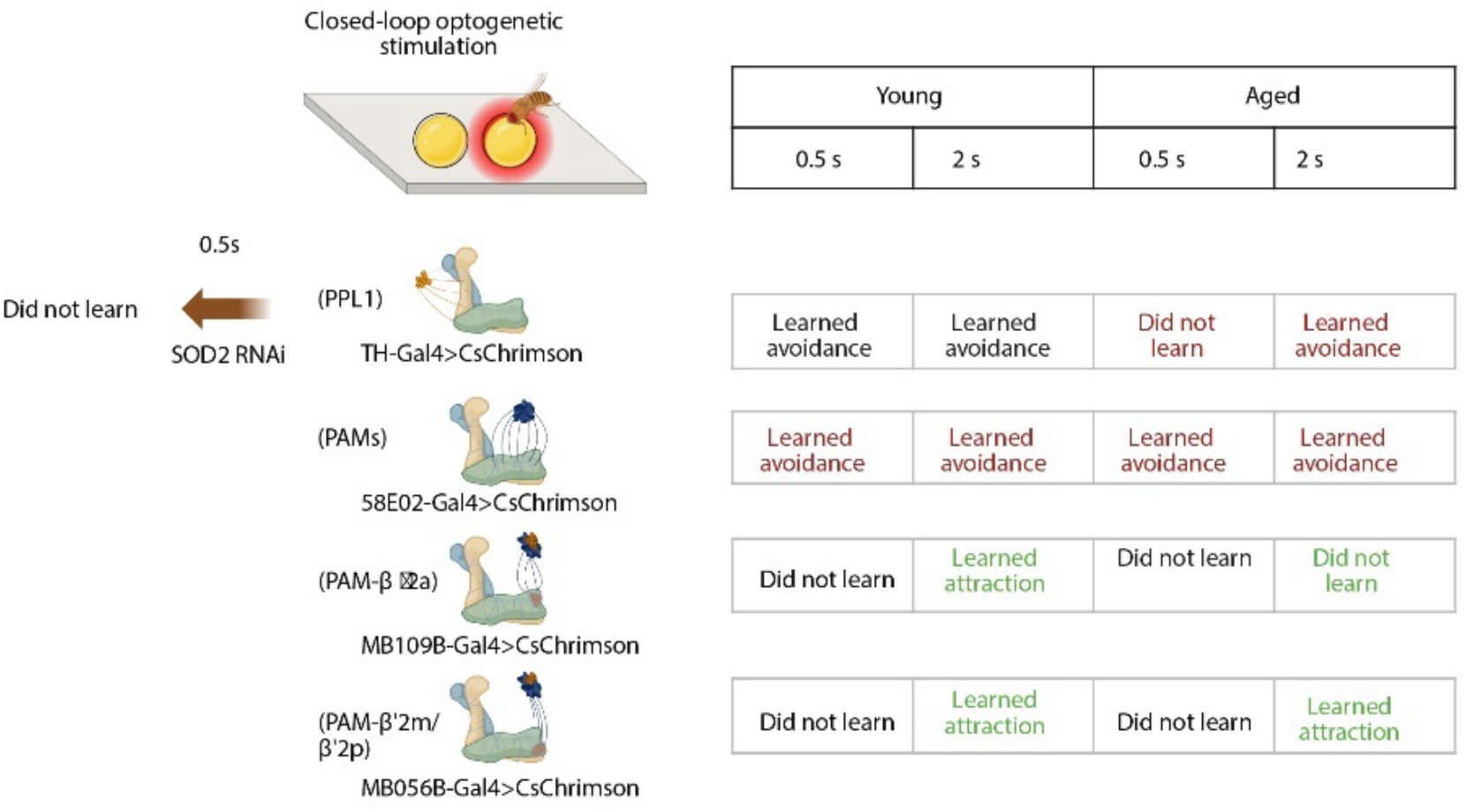
Summary of operant behavioral outcomes across dopaminergic circuits, stimulation conditions, and age. Schematic of the closed-loop optogenetic stimulation paradigm used in the optoPAD assay (left), in which sip events on the light-paired sucrose source triggered 0.5 s or 2 s red light activation of CsChrimson-expressing DANs. The table (right) summarizes the behavioral outcomes observed across genotypes, comparing young and aged flies under short (0.5 s) and prolonged (2 s) stimulation conditions. Neuronal populations include PPL1 neurons (TH-Gal4), broad PAM neurons (R58E02-Gal4), and PAM subpopulations (MB109B-Gal4; PAM-β′2a and MB056B-Gal4; PAM-β’2m/β’2p), as well as SOD2 knockdown in PPL1 neurons. Behavioral responses are color-coded as follows: learned avoidance (red), learned attraction (green), and no learning (black).

More specifically, exogenous TH-Gal4 positive DAN activation revealed that aging does not eliminate avoidance learning capacity but reduces the animal’s sensitivity to dopaminergic reinforcement. This deficit was overcome by extending the duration of optogenetic activation (Fig. 1). This indicates that MB circuits of aged flies remain capable of supporting avoidance learning when the reinforcement signal is sufficiently salient, and that the apparent learning deficit under brief stimulation reflects a reduced effectiveness of the DAN reinforcement signal rather than an inability to induce learning-related changes to the downstream neurons, such as the MBONs. Such reduced reinforcement efficacy in aged flies is consistent with evidence that aging preferentially impairs working memory-like and intermediate-term memory processes in *Drosophila* (Tamura et al., 2003; Tonoki & Davis, 2012). Because dopaminergic input from PPL1 neurons drives synaptic plasticity at Kenyon cell-MBON connections that encodes the learned association (Hige et al., 2015), a weaker or less effective reinforcement signal in aged flies would reduce the plasticity available to form the memory and produce no apparent learning even when the downstream circuitry remains relatively intact. This is consistent with our observation that strengthening reinforcement (longer stimulation) restores avoidance learning in aged animals. Notably, however, our longer stimulation paradigm also used a lower sucrose concentration to prevent premature satiation (0.5 s and 5% sucrose vs. 2 s and 2% sucrose), meaning the rescue might not be due to DAN stimulation duration alone, but also to the value of the food source. Feeding parameters analysis further showed that shorter reinforcement signals of 0.5 s were insufficient to drive learning in aged flies, whereas longer reinforcement sufficed to drive avoidance learning (though it did not fully rescue finer modulation of feeding behavior) (Fig. S1A-D).

The reason for the neuronal subtype-dependent reduction in efficacy of dopaminergic reinforcement might lie in the differential sensitivity of DANs to aging-related processes such as oxidative stress. In this scenario, TH-Gal4-positive PPL1 neurons, but not all the tested PAM neurons, in aged flies release less dopamine per optogenetic stimulation, meaning the reinforcement signal reaching the MB is too weak to reliably alter behavior. The reduced soma size observed in PPL1 but not PAM neurons in aged flies might reflect underlying changes in neuronal function such as reduced dopamine release (Fig. 5C). Alternatively, postsynaptic changes in aged MB circuits, such as reduced KC-MBON synaptic plasticity or altered dopamine receptor expression, may also contribute, given the established role of these pathways in reinforcement learning (Aso, Hattori, et al., 2014; Aso, Sitaraman, et al., 2014; Waddell, 2013). Whether either mechanism is directly altered by aging remains to be tested.

Because aging is strongly associated with mitochondrial dysfunction, we asked whether mitochondrial antioxidant capacity within PPL1 neurons is required for operant learning (Fig. 2). SOD2 is a key mitochondrial antioxidant enzyme, and its loss accelerates aspects of age-related neuronal decline in *Drosophila* (Paul et al., 2007). For instance, loss of SOD2 in olfactory projection neurons results in reduced cell body size and impaired olfactory acuity (Hussain et al., 2018). Knocking down SOD2 specifically in TH-Gal4-positive neurons abolished operant avoidance in young flies, suggesting that intact mitochondrial antioxidant function is required for these neurons to generate effective reinforcement signals. Feeding microstructure analysis supported this behavioral phenotype, as SOD2 knockdown flies showed no changes in activity bout number or feeding burst duration, consistent with the loss of avoidance observed in total sip counts (Fig. S1E-F). Mechanistically, elevated mitochondrial stress may reduce the ability of PPL1 neurons to reliably signal reinforcement during interactions with the light-paired food source, weakening the reinforcement signal reaching MB circuits. DANs may be particularly vulnerable to this kind of stress, as depletion of both SOD1 and SOD2 accelerates their loss through apoptosis (Oka et al., 2015), and dopamine metabolism itself generates additional oxidative burden through quinone formation and ROS (Graham, 1978; Meiser et al., 2013). In this sense, SOD2 knockdown may phenocopy aspects of normal aging by artificially lowering reinforcement efficacy. Future experiments such as testing whether restoring SOD2 function rescues operant avoidance learning would help to directly link mitochondrial function and reinforcement capacity.

Although PAM neurons are classically associated with appetitive reinforcement in *Drosophila* (Burke et al., 2012; Liu et al., 2012), broad optogenetic activation in our assay consistently suppressed feeding behavior across both stimulation durations and age groups (Fig. 3). Feeding microstructure analysis supported this behavioral phenotype (Fig. S2), with reduced activity bout number and feeding burst duration indicating reduced engagement with the light-paired food source. This finding aligns with recent work showing that broad PAM activation can drive acute aversion across multiple behavioral assays, including feeding, locomotion, and spatial preference (Lozada-Perdomo et al., 2025), suggesting that PAM output is more context-dependent than previously appreciated. Our findings extend this observation to an operant setting. One likely explanation is that the broad R58E02-Gal4 driver labels a heterogeneous PAM population that includes functionally distinct subtypes. While many PAM neurons promote appetitive learning, PAM-γ3 neurons within the R58E02-Gal4 expression pattern signal aversive reinforcement when activated and are suppressed by sugar ingestion (Yamagata et al., 2016). Consistent with this, PAM-γ3 activation drives avoidance in a closed-loop feeding assay (Musso et al., 2019). Repeated pairing of broad PAM activation with feeding may therefore amplify this aversive component over time. PAM population activity also encodes valence in a state-dependent manner (Siju et al., 2020), and feeding state may further shift the net output of the R58E02-Gal4 population toward aversion. Whether this reflects a true operant contingency effect or repeated exposure to an acutely aversive signal cannot be resolved with the current data.

Consistent with the established role of β′2-compartment DANs in short-term appetitive memory (Huetteroth et al., 2015), prolonged activation of both PAM-β′2a and PAM-β′2m,p drove operant attraction in young flies, indicating that these subsets provide positive reinforcement under operant conditions (Fig. 4). The failure of short stimulation to drive attraction in either line indicates that a minimum reinforcement strength is required for learning, regardless of valence or age. Notably, the two subsets dissociated with age: PAM-β′2a-driven attraction was lost in aged flies, whereas PAM-β′2m,p-driven attraction was preserved — arguing against a general age-related impairment in appetitive operant learning and instead pointing to a circuit-specific decline in dopaminergic reinforcement. Feeding parameter effects were limited and did not consistently mirror the preference phenotype (Fig. S3, S4).

Confocal imaging of PPL1 and PAM DANs (Fig. 5A, B) revealed that aging did not significantly alter the number of either population (Fig. 5C, D), indicating that age-related impairments in operant reinforcement learning are unlikely to result from significant neuronal loss (White et al., 2010). Despite this preserved cell number, PPL1 neurons showed a significant reduction in cell body surface area in aged flies (Fig. 5C), a change that may reflect altered neuronal health or metabolic state (Mattson & Magnus, 2006), as we previously observed for olfactory projection neurons under aging and oxidative stress (Hussain et al., 2018). In contrast, PAM neuron cell body size was unchanged with age (Fig. 5D), though some PAM subsets may show greater age-dependent changes than others, in line with the differential learning effects observed.

## Limitations of the study

By making reinforcement contingent on the fly’s own feeding behavior, the closed-loop design used in our study revealed age-dependent differences in reinforcement requirements that may be less apparent in paradigms where reinforcement is delivered independently of the animal’s actions. Several limitations should be noted. First, all experiments relied on optogenetic activation of defined DAN populations, which may not fully recapitulate endogenous dopamine dynamics. Second, the TH-Gal4 driver used to target PPL1 neurons also labels additional dopaminergic and some non-dopaminergic cells. Third, the age range examined here represents a moderate aging window, and more extreme ages may reveal additional impairments. Finally, the precise mechanisms by which SOD2 knockdown disrupts reinforcement signaling remain unresolved.

## Conclusions

Our results show that operant reinforcement learning in *Drosophila* depends on functionally distinct dopaminergic circuits that differ in their vulnerability to aging. Rather than causing a uniform decline in learning capacity, aging selectively reduces the strength of dopaminergic reinforcement signals, with consequences for working memory-like processes that depend on integrating recent action outcomes.

Our finding that PPL1 DANs are functionally more vulnerable than PAM neurons during operant aging challenges a purely anatomy-based view of neuronal susceptibility. While large axonal arbors are often considered a primary determinant of metabolic burden leading to functional decline, our data might point to additional parameters such as the temporal structure of neuronal activity. In operant paradigms, PPL1 neurons might be engaged in a more sustained manner, reflecting persistent negative prediction signals required for sustained avoidance. This contrasts with PAM neurons, which predominantly signal reward in a more phasic, event-linked fashion. We speculate that this difference results in a higher time-integrated calcium and metabolic load in PPL1 neurons, effectively outweighing their smaller anatomy. In this framework, vulnerability is best described by the interaction between activity and cellular scale, rather than either factor alone. Consistent with this view, sustained activity has been linked to increased mitochondrial stress (Pacelli et al., 2015; Zampese et al., 2022), reactive oxygen species production (Oswald et al., 2018), and impaired proteostasis, processes that are sensitive to disruptions in pathways such as PINK1 and Parkin (Clark et al., 2006; Ge et al., 2020; Park et al., 2006). Thus, our results support a model in which certain dopaminergic circuits are more vulnerable due to their higher duty cycle of engagement, providing a circuit-level explanation for differential aging decline within the dopaminergic system. This decline, importantly, can be compensated by prolonged reinforcement as the underlying plasticity mechanisms remain capable of learning.

## Author contributions

I.C.G.K., J.-F.D.B. and I.J. conceptualized the study and designed the experiments. I.J. carried out and analyzed most behavioral experiments, genetic manipulations and histological experiments. H.H. performed part of the optogenetic experiments and confocal imaging. H.H., J.-F.D.B., I.J. and I.C.G.K. wrote the manuscript with the help of all authors.

## Acknowledgments

We thank Natalie Lindenberg, Utsab Majumder, Martina Canova and Meenakshi for help in general fly husbandry. We wish to thank Yoshinori Aso for providing split-GAL4 lines. We acknowledge Prishita Sharma for helping in CS experiments. This project was generously funded by the German Research Foundation (GR4310/11-1 to IGK; INST 217/1135-1 to IGK) and the iBehave network, funded by the state of North Rhine-Westphalia (to IGK).

## Supplementary Materials

Figures S1 to S4

## Materials and Methods

### Data availability

All original data can be requested from the corresponding author at any time and will be made available prior to publication of this manuscript.

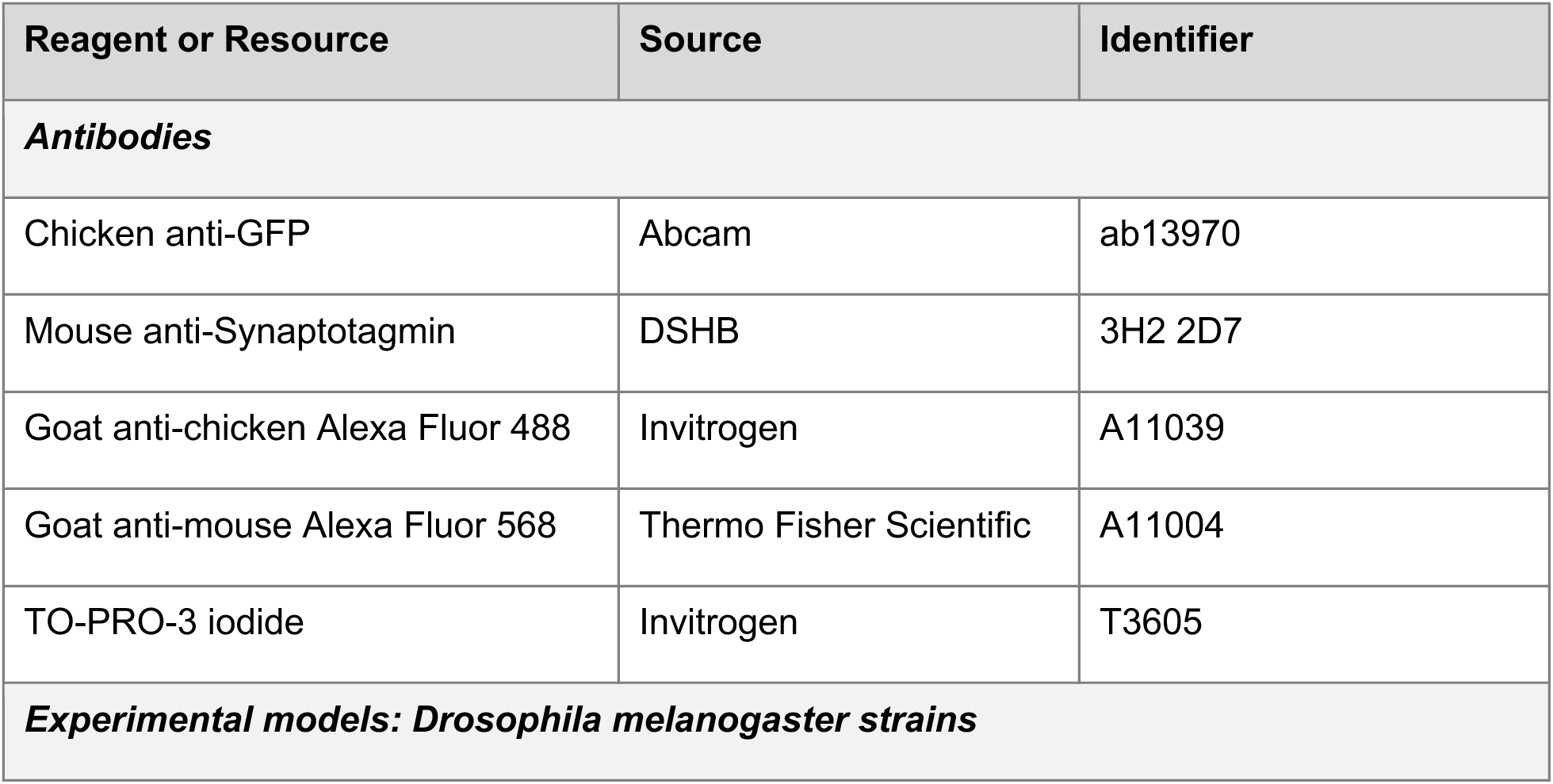
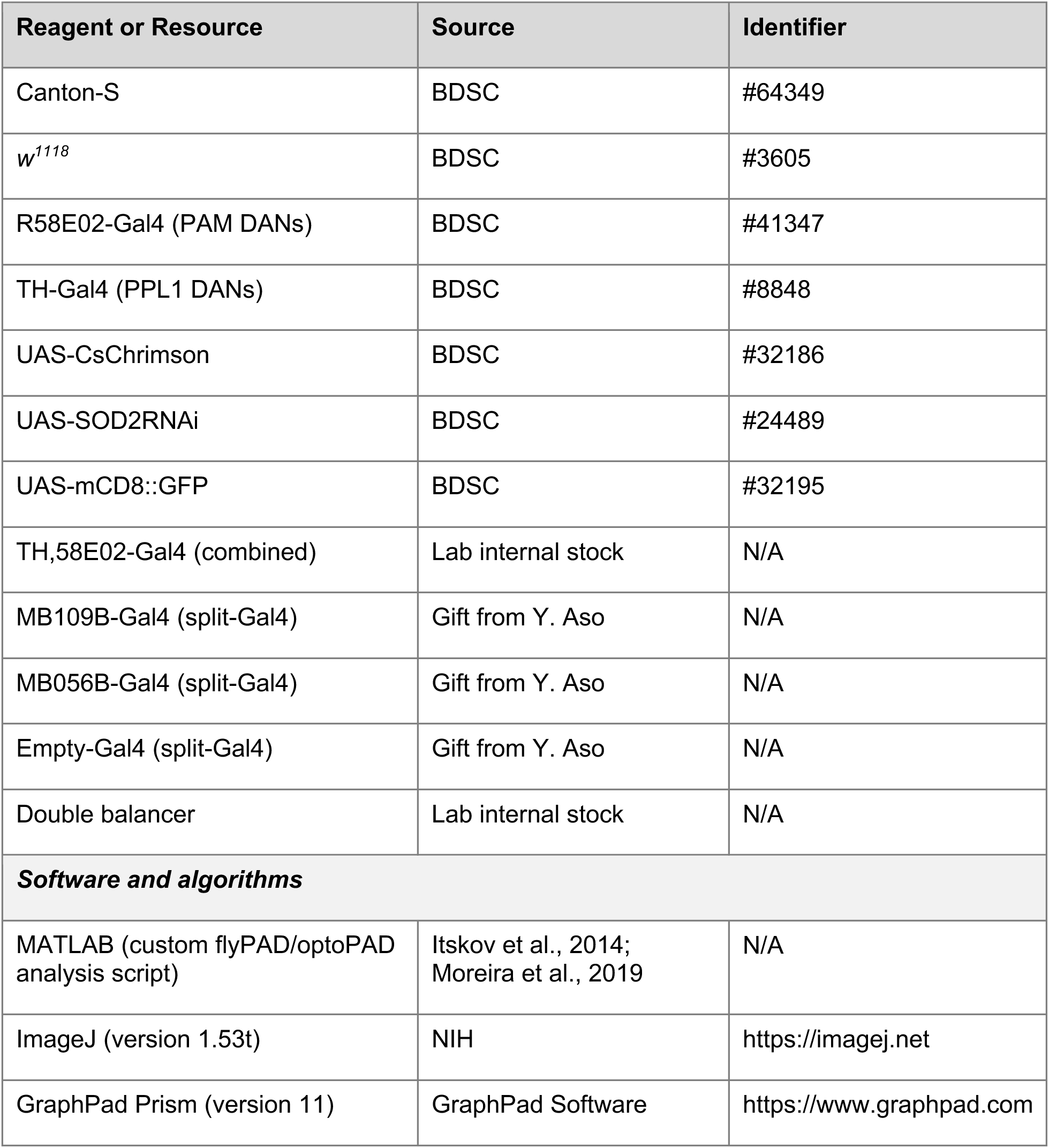

### Fly strains and maintenance

Flies were raised on standard cornmeal medium at 25°C and 60% humidity under a 12 h light/12 h dark cycle. All behavioral experiments were performed using 7–8-day-old (young) and 28–30-day-old (aged) female *Drosophila melanogaster* unless stated otherwise. For optogenetic experiments, young flies were collected 3 days after eclosion and aged flies at 25 days post-eclosion; both were transferred to food supplemented with all-trans-retinal (1:500) and maintained in darkness for 3 days. Flies were then subjected to wet starvation (water only) for 24 h before experiments. For TH-Gal4 and R58E02-Gal4 experiments, two control genotypes were used: TH-Gal4/+ or R58E02-Gal4/+ (driver-only controls) and Empty-Gal4 > UAS-CsChrimson (to control for non-specific Gal4/UAS effects). For MB109B and MB056B split-Gal4 experiments, w1118 > UAS-CsChrimson was used as the control because matched empty split-Gal4 lines were not available. All control flies were maintained on retinal-supplemented food and handled identically to experimental flies. Genotypes of all fly lines used in this study are listed in the Key Resources Table.

### flyPAD feeding assay

flyPAD experiments were conducted as described previously (Itskov et al., 2014). Flies were starved as described above prior to behavioral experiments. Individual flies were transferred by aspiration into a behavioral arena containing two food wells connected to independent electrodes. Each well contained 1% agarose (low-gelling-temperature, A9414, Sigma-Aldrich) supplemented with 5% sucrose. Experiments were performed in a climate-controlled chamber at 25°C and 60% humidity. Following transfer into the behavioral arena, feeding behavior was recorded for a fixed duration of 1 h. Capacitance signals were analyzed using a custom MATLAB script described previously. Individual feeding events (sips) were identified from capacitance changes, and feeding behavior was quantified by calculating the cumulative and total number of sips detected across the two electrodes. The same experimental procedure was used for both young and aged flies.

### optoPAD feeding assay

optoPAD experiments were conducted using the optoPAD system as previously described (Itskov et al., 2014; Moreira et al., 2019). Individual flies were transferred by aspiration into a behavioral arena containing two food wells connected to independent electrodes. Flies were offered 5% sucrose supplemented with 1% agarose (low-gelling-temperature agarose, A9414, Sigma-Aldrich). 2.5 µL of the sucrose–agarose substrate was loaded into each well prior to the start of the experiment. For closed-loop optogenetic experiments, the flyPAD system was integrated with the optoPAD platform, allowing interaction events on the designated light-paired food source to trigger red-light stimulation in real time. One food source was paired with red LED illumination (2.5 V, 625 nm); LED activation occurred upon contact with the light-paired food and remained active for a fixed duration independent of subsequent fly behavior. The light-paired food source was pseudo-randomly assigned across experiments to balance left and right positions.

Two stimulation paradigms were tested: short stimulation (0.5 s LED activation per sip event, paired with 5% sucrose) and long stimulation (2 s LED activation per sip event, paired with 2% sucrose). The lower sucrose concentration in the long-stimulation paradigm was selected to maintain comparable engagement with the food source across stimulation conditions, since longer optogenetic stimulation per sip can reduce overall feeding interactions and limit the number of reinforcement events available for learning. Feeding behavior was recorded for 1 h in a climate-controlled chamber at 25°C and 60% humidity with minimal external light. Capacitance signals were recorded and analyzed offline using a custom MATLAB script as previously described (Moreira et al., 2019). Individual feeding events (sips) were identified from capacitance changes, and feeding behavior was quantified by calculating the total and cumulative number of sips detected per electrode for each genotype. The same experimental protocol was applied to both young and aged flies.

### Immunohistochemistry and confocal imaging

Brains from young and aged flies were dissected in 0.1% PBT (PBS containing 0.1% Triton X-100) and fixed in 4% paraformaldehyde in PBS for 40 min at room temperature. Samples were washed twice for 10 min in 0.1% PBT followed by one wash in PBS for 10 min, then blocked for 30 min in 10% normal goat serum in 0.1% PBT. Primary antibodies were diluted in blocking solution and incubated overnight at 4°C. The following primary antibodies were used: chicken anti-GFP (1:1000; Abcam, ab13970) and mouse anti-synaptotagmin (1:50; DSHB, 3H2 2D7).

The following day, samples were washed three times for 20 min each in 0.1% PBT at 4°C. Secondary antibodies (goat anti-chicken Alexa Fluor 488, 1:200, Invitrogen A11039; goat anti-mouse Alexa Fluor 568, 1:200, Thermo Fisher Scientific A11004) were diluted in blocking solution and incubated overnight at 4°C, together with TO-PRO-3 iodide (1:200; Invitrogen, T3605) for nuclear counterstaining. The next morning, brains were washed once for 20 min and once for 1 h in 0.1% PBT at 4°C, then mounted on glass slides under coverslips using Vectashield mounting medium (Vector Laboratories). Images were acquired sequentially on a Leica SP5 confocal microscope with a 20× water immersion objective (1024 × 1024 pixels), using the 488 nm laser line. Z-stacks were acquired with a step size of 1 µm and line averaging of 4. PAM neurons were imaged at zoom factor 3.2 with 10% laser power; PPL1 neurons at zoom factor 8 with 15% laser power. All samples within an experimental set were acquired with identical settings. Image data were processed and analyzed using ImageJ (version 1.53t). Although synaptotagmin staining was acquired during imaging, it did not produce reliable signal under the experimental conditions used and was therefore excluded from downstream analyses.

Neuron cell bodies were identified based on GFP signal intensity and anatomical position and manually counted throughout the complete z-stack for each brain hemisphere. Maximum intensity projections were used for visualization only; quantification was performed on the full z-stack. Counts from the left and right hemispheres were averaged to obtain a single value per brain. Neuron cell body surface area was quantified by manually outlining individual somata using the polygon selection tool on the optical section showing the maximum cross-sectional area of each soma. Areas (µm²) were measured using the ‘Measure’ function with the scale calibrated from confocal metadata. To anchor measurements against published anatomy, PPL1 soma sizes were also measured from confocal stacks of the FlyLight split-GAL4 line MB320C using the same protocol, yielding values comparable to those obtained from our own imaging.

### Quantification and statistical analysis

Feeding behavior was quantified using the following parameters as previously defined (Itskov et al., 2014; Moreira et al., 2019): total sip number (number of feeding events detected at 60 min), cumulative sip number (feeding events accumulated across the assay), activity bout number (discrete feeding engagement periods separated by inactivity), total activity bout duration (summed duration of activity bouts), and feeding burst duration (duration of continuous feeding bursts within an activity bout).

Flies were included in the analysis only if control genotypes (Gal4/+, UAS/+ and UAS-SOD2i/+) showed no systematic preference between light-paired and non-light-paired food sources, confirming the absence of inherent positional bias. Recordings with fewer than 10 total sips were excluded. The sample size (n) in all experiments denotes the number of flies. Each experiment was carried out at least twice independently with different parental crosses on different days to ensure repeatability. Controls and test groups were run side-by-side, and the order and position of optoPAD arenas were randomized.

All statistical analyses were performed using GraphPad Prism (version 11). Paired comparisons between light-paired and non-light-paired food sources within the same fly were analyzed using the Wilcoxon matched-pairs signed-rank test as indicated in the figure legends. Light OFF (blue; grey) and light ON (orange; purple). Cumulative sips over time plots are presented as mean ± SEM. Boxplots represent the medians, 25th and 75th percentiles of the data distributions. Whiskers are plotted according to Tukey: 25th percentile – 1.5*IQR and 75th + 1.5*IQR. Values outside this range are plotted as single data points. Baseline comparisons between young and aged WT-CS (Fig. 1A) flies were analyzed using the Mann–Whitney U test. Statistical notation: ns, p > 0.05; * p < 0.05; ** p < 0.01; *** p < 0.001; **** p < 0.0001.

## References

1. Aso, Y., Hattori, D., Yu, Y., Johnston, R. M., Iyer, N. A., Ngo, T.-T., Dionne, H., Abbott, L. F., Axel, R., Tanimoto, H., & Rubin, G. M. (2014). The neuronal architecture of the mushroom body provides a logic for associative learning. eLife, 3, e04577. 10.7554/eLife.04577

2. Aso, Y., Sitaraman, D., Ichinose, T., Kaun, K. R., Vogt, K., Belliart-Guérin, G., Plaçais, P.-Y., Robie, A. A., Yamagata, N., Schnaitmann, C., Rowell, W. J., Johnston, R. M., Ngo, T.-T. B., Chen, N., Korff, W., Nitabach, M. N., Heberlein, U., Preat, T., Branson, K. M., … Rubin, G. M. (2014). Mushroom body output neurons encode valence and guide memory-based action selection in Drosophila. eLife, 3. 10.7554/eLife.04580

3. Aso, Y., Siwanowicz, I., Bräcker, L., Ito, K., Kitamoto, T., & Tanimoto, H. (2010). Specific dopaminergic neurons for the formation of labile aversive memory. Curr Biol, 20(16), 1445–1451. 10.1016/j.cub.2010.06.048

4. Bäckman, L., Nyberg, L., Lindenberger, U., Li, S.-C., & Farde, L. (2006). The correlative triad among aging, dopamine, and cognition: Current status and future prospects. Neuroscience & Biobehavioral Reviews, 30(6), 791–807. 10.1016/j.neubiorev.2006.06.005

5. Blazer, D. G., Yaffe, K., Liverman, C. T., Aging, C. on the P. H. D. of C., Policy, B. on H. S., & Medicine, I. of. (2015). Cognitive Aging: Progress in Understanding and Opportunities for Action (D. G. Blazer, K. Yaffe, & C. T. Liverman, Eds.). National Academies Press (US). 10.17226/21693

6. Brooker, S. M., Naylor, G. E., & Krainc, D. (2024). Cell biology of Parkinson’s disease: Mechanisms of synaptic, lysosomal, and mitochondrial dysfunction. Current Opinion in Neurobiology, 85, 102841. 10.1016/j.conb.2024.102841

7. Burbulla, L. F., Song, P., Mazzulli, J. R., Zampese, E., Wong, Y. C., Jeon, S., Santos, D. P., Blanz, J., Obermaier, C. D., Strojny, C., Savas, J. N., Kiskinis, E., Zhuang, X., Krüger, R., Surmeier, D. J., & Krainc, D. (2017). Dopamine oxidation mediates mitochondrial and lysosomal dysfunction in Parkinson’s disease. Science, 357(6357), 1255–1261. 10.1126/science.aam9080

8. Burke, C. J., Huetteroth, W., Owald, D., Perisse, E., Krashes, M. J., Das, G., Gohl, D., Silies, M., Certel, S., & Waddell, S. (2012). Layered reward signalling through octopamine and dopamine in *Drosophila*. Nature, 492(7429), 433–437. 10.1038/nature11614

9. Clark, I. E., Dodson, M. W., Jiang, C., Cao, J. H., Huh, J. R., Seol, J. H., Yoo, S. J., Hay, B. A., & Guo, M. (2006). *Drosophila* pink1 is required for mitochondrial function and interacts genetically with parkin. Nature, 441(7097), 1162–1166. 10.1038/nature04779

10. Coleman, C. R., Pallos, J., Arreola-Bustos, A., Wang, L., Raftery, D., Promislow, D. E. L., & Martin, I. (2024). Natural variation in age-related dopamine neuron degeneration is glutathione dependent and linked to life span. Proceedings of the National Academy of Sciences, 121(42). 10.1073/pnas.2403450121

11. Davie, K., Janssens, J., Koldere, D., De Waegeneer, M., Pech, U., Kreft, Ł., Aibar, S., Makhzami, S., Christiaens, V., Bravo González-Blas, C., Poovathingal, S., Hulselmans, G., Spanier, K. I., Moerman, T., Vanspauwen, B., Geurs, S., Voet, T., Lammertyn, J., Thienpont, B., … Aerts, S. (2018). A Single-Cell Transcriptome Atlas of the Aging *Drosophila* Brain. Cell, 174(4), 982–998.e20. 10.1016/j.cell.2018.05.057

12. Friggi-Grelin, F., Coulom, H., Meller, M., Gomez, D., Hirsh, J., & Birman, S. (2003). Targeted gene expression in *Drosophila* dopaminergic cells using regulatory sequences from tyrosine hydroxylase. Journal of Neurobiology, 54(4), 618–627. 10.1002/neu.10185

13. Galili, D. S., Dylla, K. V., Lüdke, A., Friedrich, A. B., Yamagata, N., Wong, J. Y. H., Ho, C. H., Szyszka, P., & Tanimoto, H. (2014). Converging Circuits Mediate Temperature and Shock Aversive Olfactory Conditioning in *Drosophila*. Current Biology, 24(15), 1712–1722. 10.1016/j.cub.2014.06.062

14. Ge, P., Dawson, V. L., & Dawson, T. M. (2020). PINK1 and Parkin mitochondrial quality control: a source of regional vulnerability in Parkinson’s disease. Molecular Neurodegeneration, 15(1), 20. 10.1186/s13024-020-00367-7

15. Graham, D. G. (1978). Oxidative pathways for catecholamines in the genesis of neuromelanin and cytotoxic quinones. Molecular Pharmacology, 14(4), 633–643.

16. Greene, J. C., Whitworth, A. J., Kuo, I., Andrews, L. A., Feany, M. B., & Pallanck, L. J. (2003). Mitochondrial pathology and apoptotic muscle degeneration in *Drosophila* parkin mutants. Proceedings of the National Academy of Sciences of the United States of America, 100(7), 4078–4083. 10.1073/pnas.0737556100

17. Griffith, L. C. (2012). Identifying behavioral circuits in *Drosophila melanogaster*: moving targets in a flying insect. Current Opinion in Neurobiology, 22(4), 609–614. 10.1016/j.conb.2012.01.002

18. Hige, T., Aso, Y., Modi, M. N., Rubin, G. M., & Turner, G. C. (2015). Heterosynaptic Plasticity Underlies Aversive Olfactory Learning in *Drosophila*. Neuron, 88(5), 985–998. 10.1016/j.neuron.2015.11.003

19. Hirth, F. (2010). *Drosophila melanogaster* in the study of human neurodegeneration. CNS & Neurological Disorders Drug Targets, 9(4), 504–523. 10.2174/187152710791556104

20. Huetteroth, W., Perisse, E., Lin, S., Klappenbach, M., Burke, C., & Waddell, S. (2015). Sweet Taste and Nutrient Value Subdivide Rewarding Dopaminergic Neurons in *Drosophila*. Current Biology, 25(6), 751–758. 10.1016/j.cub.2015.01.036

21. Hussain, A., Pooryasin, A., Zhang, M., Loschek, L. F., La Fortezza, M., Friedrich, A. B., Blais, C.-M., Üçpunar, H. K., Yépez, V. A., Lehmann, M., Gompel, N., Gagneur, J., Sigrist, S. J., & Grunwald Kadow, I. C. (2018). Inhibition of oxidative stress in cholinergic projection neurons fully rescues aging-associated olfactory circuit degeneration in *Drosophila*. eLife, 7. 10.7554/eLife.32018

22. Itskov, P. M., Moreira, J. M., Vinnik, E., Lopes, G., Safarik, S., Dickinson, M. H., & Ribeiro, C. (2014). Automated monitoring and quantitative analysis of feeding behaviour in *Drosophila*. Nat Commun, 5, 4560. 10.1038/ncomms5560

23. Klapoetke, N. C., Murata, Y., Kim, S. S., Pulver, S. R., Birdsey-Benson, A., Cho, Y. K., Morimoto, T. K., Chuong, A. S., Carpenter, E. J., Tian, Z., Wang, J., Xie, Y., Yan, Z., Zhang, Y., Chow, B. Y., Surek, B., Melkonian, M., Jayaraman, V., Constantine-Paton, M., … Boyden, E. S. (2014). Independent optical excitation of distinct neural populations. Nature Methods, 11(3), 338–346. 10.1038/nmeth.2836

24. Lewis, L. P. C., Siju, K. P., Aso, Y., Friedrich, A. B., Bulteel, A. J. B., Rubin, G. M., & Grunwald Kadow, I. C. (2015). A Higher Brain Circuit for Immediate Integration of Conflicting Sensory Information in *Drosophila*. Current Biology, 25(17), 2203–2214. 10.1016/j.cub.2015.07.015

25. Liu, C., Plaçais, P. Y., Yamagata, N., Pfeiffer, B. D., Aso, Y., Friedrich, A. B., Siwanowicz, I., Rubin, G. M., Preat, T., & Tanimoto, H. (2012). A subset of dopamine neurons signals reward for odour memory in *Drosophila*. Nature, 488(7412), 512–516. 10.1038/nature11304

26. López-Otín, C., Blasco, M. A., Partridge, L., Serrano, M., & Kroemer, G. (2013). The Hallmarks of Aging. Cell, 153(6), 1194–1217. 10.1016/j.cell.2013.05.039

27. Lozada-Perdomo, F. V, Chen, Y., Jacobs, R. V, Yeo, J., Yang, M. M., Bhalerao, J., & Devineni, A. V. (2025). Dual roles of *Drosophila* reward-encoding dopamine neurons in regulating innate and learned behaviors. IScience, 28(11), 113817. 10.1016/j.isci.2025.113817

28. Mao, Z., & Davis, R. L. (2009). Eight different types of dopaminergic neurons innervate the *Drosophila* mushroom body neuropil: anatomical and physiological heterogeneity. Frontiers in Neural Circuits, 3, 5. 10.3389/neuro.04.005.2009

29. Masek, P., Worden, K., Aso, Y., Rubin, G. M., & Keene, A. C. (2015). A Dopamine-Modulated Neural Circuit Regulating Aversive Taste Memory in *Drosophila*. Current Biology, 25(11), 1535–1541. 10.1016/j.cub.2015.04.027

30. Matsuno, M., Uemura, N., Miyashita, T., Kobayashi, K. S., Liotta, A., Kimura, M., Ofusa, K., Owald, D., Matsuo, N., Saitoe, M., & Horiuchi, J. (2026). Aberrant dopaminergic activity during consolidation causes age-related memory generalization in *Drosophila*. PLOS Biology, 24(4), e3003752. 10.1371/journal.pbio.3003752

31. Mattson, M. P., & Magnus, T. (2006). Ageing and neuronal vulnerability. Nature Reviews Neuroscience, 7(4), 278–294. 10.1038/nrn1886

32. Meiser, J., Weindl, D., & Hiller, K. (2013). Complexity of dopamine metabolism. Cell Communication and Signaling, 11(1), 34. 10.1186/1478-811X-11-34

33. Moreira, J. M., Itskov, P. M., Goldschmidt, D., Baltazar, C., Steck, K., Tastekin, I., Walker, S. J., & Ribeiro, C. (2019). optoPAD, a closed-loop optogenetics system to study the circuit basis of feeding behaviors. eLife, 8, e43924. 10.7554/eLife.43924

34. Murman, D. (2015). The Impact of Age on Cognition. Seminars in Hearing, 36(03), 111–121. 10.1055/s-0035-1555115

35. Murphy, E. S., & Lupfer, G. J. (2014). Basic Principles of Operant Conditioning. In The Wiley Blackwell Handbook of Operant and Classical Conditioning (pp. 165–194). Wiley. 10.1002/9781118468135.ch8

36. Musso, P.-Y., Junca, P., Jelen, M., Feldman-Kiss, D., Zhang, H., Chan, R. C., & Gordon, M. D. (2019). Closed-loop optogenetic activation of peripheral or central neurons modulates feeding in freely moving *Drosophila*. eLife, 8. 10.7554/eLife.45636

37. Neckameyer, W. S., Woodrome, S., Holt, B., & Mayer, A. (2000). Dopamine and senescence in *Drosophila melanogaster*. Neurobiology of Aging, 21(1), 145–152. 10.1016/S0197-4580(99)00109-8

38. Oka, S., Hirai, J., Yasukawa, T., Nakahara, Y., & Inoue, Y. H. (2015). A correlation of reactive oxygen species accumulation by depletion of superoxide dismutases with age-dependent impairment in the nervous system and muscles of *Drosophila* adults. Biogerontology, 16(4), 485–501. 10.1007/s10522-015-9570-3

39. Oswald, M. C., Brooks, P. S., Zwart, M. F., Mukherjee, A., West, R. J., Giachello, C. N., Morarach, K., Baines, R. A., Sweeney, S. T., & Landgraf, M. (2018). Reactive oxygen species regulate activity-dependent neuronal plasticity in *Drosophila*. eLife, 7. 10.7554/eLife.39393

40. Pacelli, C., Giguère, N., Bourque, M.-J., Lévesque, M., Slack, R. S., & Trudeau, L.-É. (2015). Elevated Mitochondrial Bioenergetics and Axonal Arborization Size Are Key Contributors to the Vulnerability of Dopamine Neurons. Current Biology, 25(18), 2349–2360. 10.1016/j.cub.2015.07.050

41. Park, J., Lee, S. B., Lee, S., Kim, Y., Song, S., Kim, S., Bae, E., Kim, J., Shong, M., Kim, J.-M., & Chung, J. (2006). Mitochondrial dysfunction in *Drosophila* PINK1 mutants is complemented by parkin. Nature, 441(7097), 1157–1161. 10.1038/nature04788

42. Paul, A., Belton, A., Nag, S., Martin, I., Grotewiel, M. S., & Duttaroy, A. (2007). Reduced mitochondrial SOD displays mortality characteristics reminiscent of natural aging. Mechanisms of Ageing and Development, 128(11–12), 706–716. 10.1016/j.mad.2007.10.013

43. Perisse, E., Owald, D., Barnstedt, O., Talbot, C. B., Huetteroth, W., & Waddell, S. (2016). Aversive learning and appetitive motivation toggle feed-forward inhibition in the *Drosophila* mushroom body. Neuron, 90(5), 1086–1099. 10.1016/j.neuron.2016.04.034

44. Piper, M. D. W., & Partridge, L. (2018). *Drosophila* as a model for ageing. Biochimica et Biophysica Acta (BBA) - Molecular Basis of Disease, 1864(9), 2707–2717. 10.1016/j.bbadis.2017.09.016

45. Rajagopalan, A. E., Darshan, R., Hibbard, K. L., Fitzgerald, J. E., & Turner, G. C. (2023). Reward expectations direct learning and drive operant matching in *Drosophila*. Proceedings of the National Academy of Sciences, 120(39), e2221415120. 10.1073/pnas.2221415120

46. Riemensperger, T., Völler, T., Stock, P., Buchner, E., & Fiala, A. (2005). Punishment Prediction by Dopaminergic Neurons in *Drosophila*. Current Biology, 15(21), 1953–1960. 10.1016/j.cub.2005.09.042

47. Salthouse, T. A. (1994). The aging of working memory. Neuropsychology, 8(4), 535–543. 10.1037/0894-4105.8.4.535

48. Siju, K. P., Štih, V., Aimon, S., Gjorgjieva, J., Portugues, R., & Grunwald Kadow, I. C. (2020). Valence and State-Dependent Population Coding in Dopaminergic Neurons in the Fly Mushroom Body. Current Biology, 30(11), 2104–2115.e4. 10.1016/j.cub.2020.04.037

49. Tamura, T., Chiang, A.-S., Ito, N., Liu, H.-P., Horiuchi, J., Tully, T., & Saitoe, M. (2003). Aging specifically impairs amnesiac dependent memory in *Drosophila*. Neuron, 40(5), 1003–1011. 10.1016/S0896-6273(03)00732-3

50. Tonoki, A., & Davis, R. L. (2012). Aging impairs intermediate-term behavioral memory by disrupting the dorsal paired medial neuron memory trace. Proceedings of the National Academy of Sciences of the United States of America, 109(16), 6319–6324. 10.1073/pnas.1118126109

51. Tonoki, A., Ogasawara, M., Yu, Z., & Itoh, M. (2020). Appetitive Memory with Survival Benefit Is Robust Across Aging in *Drosophila*. The Journal of Neuroscience, 40(11), 2296–2304. 10.1523/JNEUROSCI.2045-19.2020

52. Trist, B. G., Hare, D. J., & Double, K. L. (2019). Oxidative stress in the aging substantia nigra and the etiology of Parkinson’s disease. Aging Cell, 18(6). 10.1111/acel.13031

53. Waddell, S. (2013). Reinforcement signaling in *Drosophila*; dopamine does it all after all. Curr Opin Neurobiol, 23(3), 324–329. 10.1016/j.conb.2012.10.005

54. Watanabe, H., Dijkstra, J. M., & Nagatsu, T. (2024). Parkinson’s Disease: Cells Succumbing to Lifelong Dopamine-Related Oxidative Stress and Other Bioenergetic Challenges. International Journal of Molecular Sciences, 25(4), 2009. 10.3390/ijms25042009

55. White, K. E., Humphrey, D. M., & Hirth, F. (2010). The dopaminergic system in the aging brain of *Drosophila*. Front Neurosci, 4, 205. 10.3389/fnins.2010.00205

56. Wiggin, T. D., Hsiao, Y., Liu, J. B., Huber, R., & Griffith, L. C. (2021). Rest Is Required to Learn an Appetitively-Reinforced Operant Task in *Drosophila*. Frontiers in Behavioral Neuroscience, Volume 15-2021. 10.3389/fnbeh.2021.681593

57. Yamagata, N., Hiroi, M., Kondo, S., Abe, A., & Tanimoto, H. (2016). Suppression of Dopamine Neurons Mediates Reward. PLoS Biology, 14(12), e1002586. 10.1371/journal.pbio.1002586

58. Yamagata, N., Ichinose, T., Aso, Y., Plaçais, P.-Y., Friedrich, A. B., Sima, R. J., Preat, T., Rubin, G. M., & Tanimoto, H. (2015). Distinct dopamine neurons mediate reward signals for short- and long-term memories. Proceedings of the National Academy of Sciences, 112(2), 578–583. 10.1073/pnas.1421930112

59. Zampese, E., Wokosin, D. L., Gonzalez-Rodriguez, P., Guzman, J. N., Tkatch, T., Kondapalli, J., Surmeier, W. C., D’Alessandro, K. B., De Stefani, D., Rizzuto, R., Iino, M., Molkentin, J. D., Chandel, N. S., Schumacker, P. T., & Surmeier, D. J. (2022). Ca2+ channels couple spiking to mitochondrial metabolism in substantia nigra dopaminergic neurons. Science Advances, 8(39). 10.1126/sciadv.abp8701

