## Supplementary Figures for "Reduced dopaminergic reinforcement, not learning capacity, limits operant learning in aging *Drosophila*"

**Supplementary Figures 1-4**

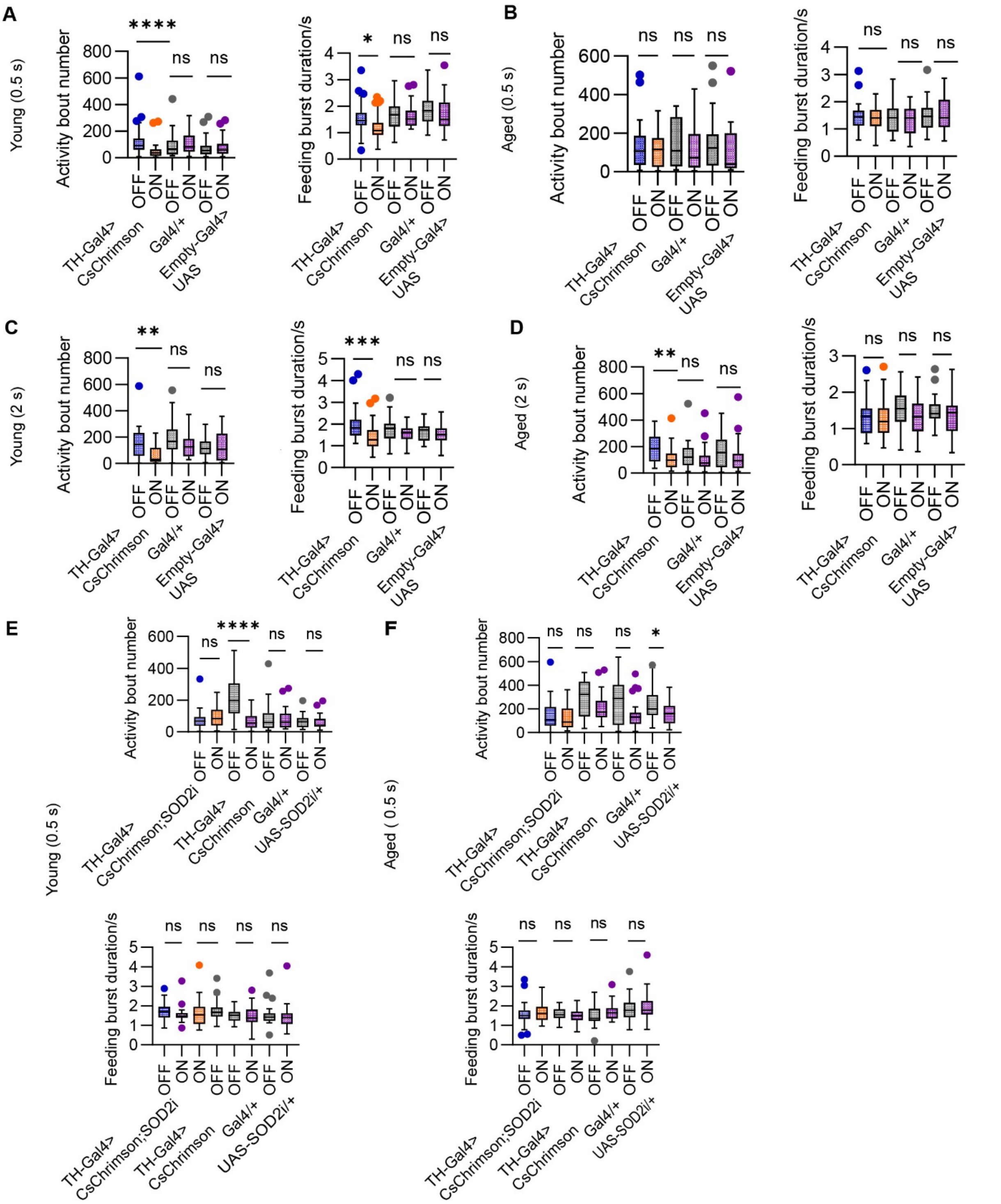


**Figure S1. Feeding parameters analysis under optogenetic activation of dopaminergic PPL1 neurons and SOD2 RNAi knockdown (Related to Figure 1 and 2)**

(A–B) Feeding parameters analysis following short optogenetic activation of dopaminergic PPL1 neurons (0.5 s, 5% sucrose) in young (A) and aged (B) flies. Activity bout number (left) and feeding burst duration (right) are shown under light OFF and light ON conditions for TH-Gal4>CsChrimson experimental flies (young n = 31, aged n = 29), Gal4/+ controls (young n = 29, aged n = 30), and Empty-Gal4>UAS controls (young n = 30, aged n = 29).

(C–D) Feeding parameters analysis following prolonged optogenetic activation of dopaminergic PPL1 neurons (2 s, 2% sucrose) in young (C) and aged (D) flies. Activity bout number (left) and feeding burst duration (right) are shown under light OFF and light ON conditions for TH-Gal4>CsChrimson experimental flies (young n = 29, aged n = 29), Gal4/+ controls (young n = 28, aged n = 30), and Empty-Gal4>UAS controls (young n = 31, aged n = 29).

(E–F) Feeding parameters analysis following optogenetic activation of dopaminergic PPL1 neurons under SOD2 knockdown conditions (0.5 s stimulation) in young (E) and aged (F) flies. Activity bout number (top) and feeding burst duration (bottom) are shown for TH-Gal4>UAS-CsChrimson; SOD2i flies (young n = 21, aged n = 25), TH-Gal4>CsChrimson flies (young n = 21, aged n = 22), Gal4/+ controls (young n = 21, aged n = 24) and UAS-SOD2i/+ controls (young n = 22, aged n = 28) under light OFF and light ON conditions.

Light OFF (blue; grey) and light ON (orange; purple). Statistical comparisons were performed using the Wilcoxon matched-pairs signed-rank test. (A) Young (0.5 s) Activity bout number: TH>CsChrimson, $p < 0.0001$; Gal4/+, $p = 0.3223$; Empty-Gal4>UAS, $p = 0.6739$. Feeding burst duration: TH>CsChrimson, $p = 0.0474$; Gal4/+, $p = 0.9330$; Empty-Gal4>UAS, $p = 0.3818$. (B) Aged (0.5 s), Activity bout number: TH>CsChrimson, $p = 0.9703$; Gal4/+, $p = 0.2917$; Empty-Gal4>UAS, $p = 0.1067$. Feeding burst duration: TH>CsChrimson, $p = 0.2193$; Gal4/+, $p = 0.6408$; Empty-Gal4>UAS, $p = 0.7836$. (C) Young (2 s), Activity bout number: TH>CsChrimson, $p = 0.0024$; Gal4/+, $p = 0.1987$; Empty-Gal4>UAS, $p = 0.9889$. Feeding burst duration: TH>CsChrimson, $p = 0.0002$; Gal4/+, $p = 0.1105$; Empty-Gal4>UAS, $p = 0.4149$. (D) Aged (2 s), Activity bout number: TH>CsChrimson, $p = 0.0079$; Gal4/+, $p = 0.1474$; Empty-Gal4>UAS, $p = 0.1167$. Feeding burst duration: TH>CsChrimson, $p = 0.1017$; Gal4/+, $p = 0.1106$; Empty-Gal4>UAS, $p = 0.1775$. (E) Young (0.5 s), Activity bout number: TH>CsChrimson;SOD2i, $p = 0.2323$; TH>CsChrimson, $p < 0.0001$; Gal4/+, $p = 0.7335$; UAS-SOD2i/+, $p = 0.6200$. Feeding burst duration: TH>CsChrimson;SOD2i, $p = 0.2064$; TH>CsChrimson, $p = 0.3784$; Gal4/+, $p = 0.8695$; UAS-SOD2i/+, $p = 0.4028$. (F) Aged (0.5 s), Activity bout number: TH>CsChrimson;SOD2i, $p = 0.4865$; TH>CsChrimson, $p = 0.2029$; Gal4/+, $p = 0.1090$; UAS-SOD2i/+, $p = 0.0432$. Feeding burst duration: TH>CsChrimson;SOD2i, $p = 0.6282$; TH>CsChrimson, $p = 0.1266$; Gal4/+, $p = 0.1875$; UAS-SOD2i/+, $p = 0.3779$.

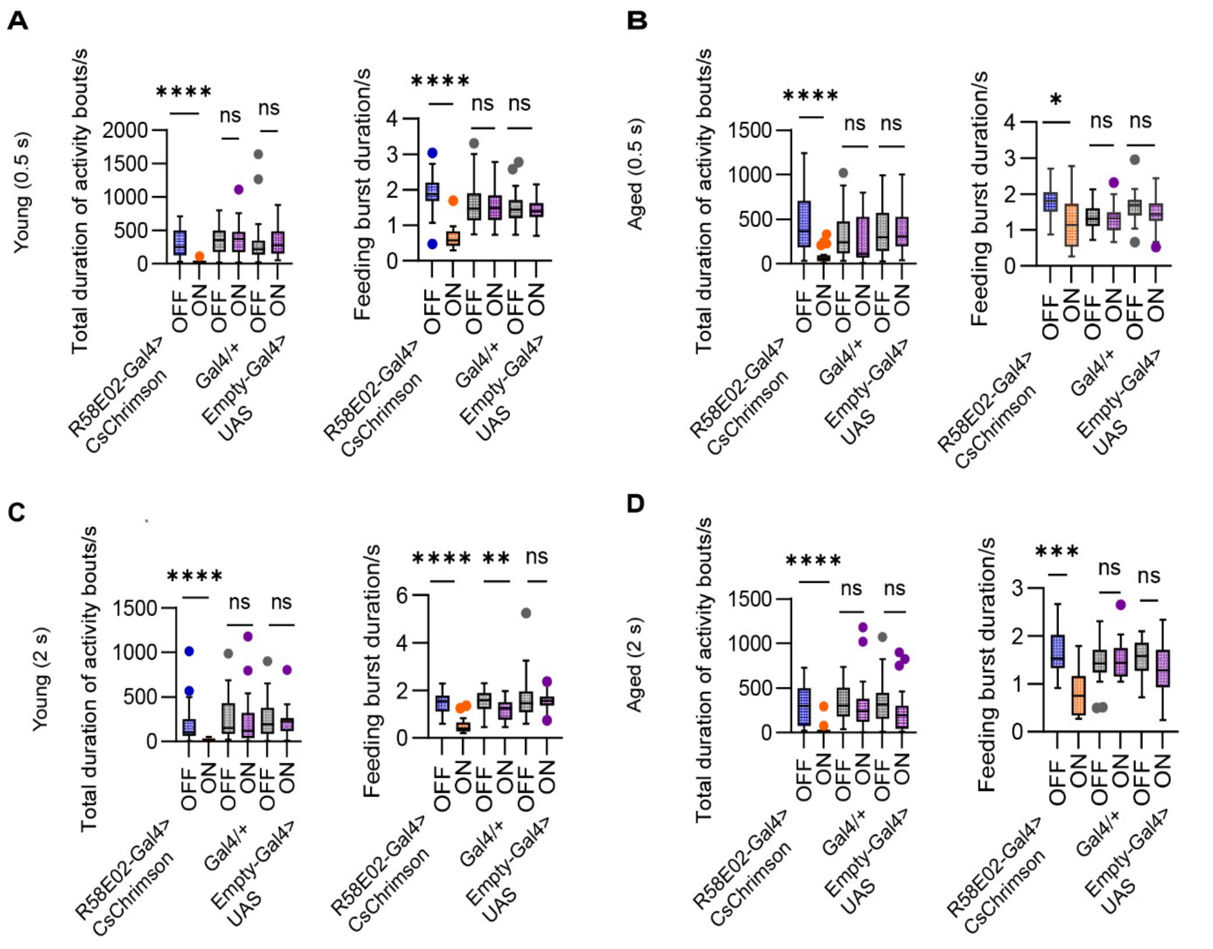


**Figure S2. Feeding parameters analysis following activation of dopaminergic PAM neurons (Related to Figure 3)**

(A–B) Feeding parameters analysis following short optogenetic activation (0.5 s, 5% sucrose) of dopaminergic PAM neurons in young (A) and aged (B) flies. Total duration of activity bouts (left) and feeding burst duration (right) are shown under light OFF and light ON conditions for R58E02-Gal4>CsChrimson experimental flies (young n = 25, aged n = 22), Gal4/+ controls (young n = 29, aged n = 30), and Empty-Gal4>UAS controls (young n = 30, aged n = 28).

(C–D) Feeding parameters analysis following prolonged optogenetic activation (2 s, 2% sucrose) of dopaminergic PAM neurons in young (C) and aged (D) flies. Total duration of activity bouts (left) and feeding burst duration (right) are shown under light OFF and light ON conditions for R58E02-Gal4>CsChrimson experimental flies (young n = 28, aged n = 21), Gal4/+ controls (young n = 32, aged n = 30), and Empty-Gal4>UAS controls (young n = 27, aged n = 27).

Light OFF (blue; grey) and light ON (orange; purple). Statistical comparisons were performed using the Wilcoxon matched-pairs signed-rank test. (A) Young (0.5 s), Total duration of activity bouts: R58E02>CsChrimson, $p < 0.0001$; Gal4/+, $p = 0.6856$; Empty-Gal4>UAS, $p = 0.4280$. Feeding burst duration: R58E02>CsChrimson, $p < 0.0001$; Gal4/+, $p = 0.5048$; Empty-Gal4>UAS, $p = 0.4614$. (B) Aged (0.5 s), Total duration of activity bouts: R58E02>CsChrimson, $p < 0.0001$; Gal4/+, $p = 0.5425$; Empty-Gal4>UAS, $p = 0.9911$. Feeding burst duration: R58E02>CsChrimson, $p = 0.0117$; Gal4/+, $p = 0.8437$; Empty-Gal4>UAS, $p = 0.5950$. (C) Young (2 s), Total duration of activity bouts: R58E02>CsChrimson, $p < 0.0001$; Gal4/+, $p = 0.0803$; Empty-Gal4>UAS, $p = 0.8223$. Feeding burst duration: R58E02>CsChrimson, $p < 0.0001$; Gal4/+, $p = 0.0025$; Empty-Gal4>UAS, $p = 0.9765$. (D) Aged (2 s), Total duration of activity bouts: R58E02>CsChrimson, $p < 0.0001$; Gal4/+, $p = 0.2286$; Empty-Gal4>UAS, $p = 0.1167$. Feeding burst duration: R58E02>CsChrimson, $p = 0.0001$; Gal4/+, $p = 0.8621$; Empty-Gal4>UAS, $p = 0.2437$.

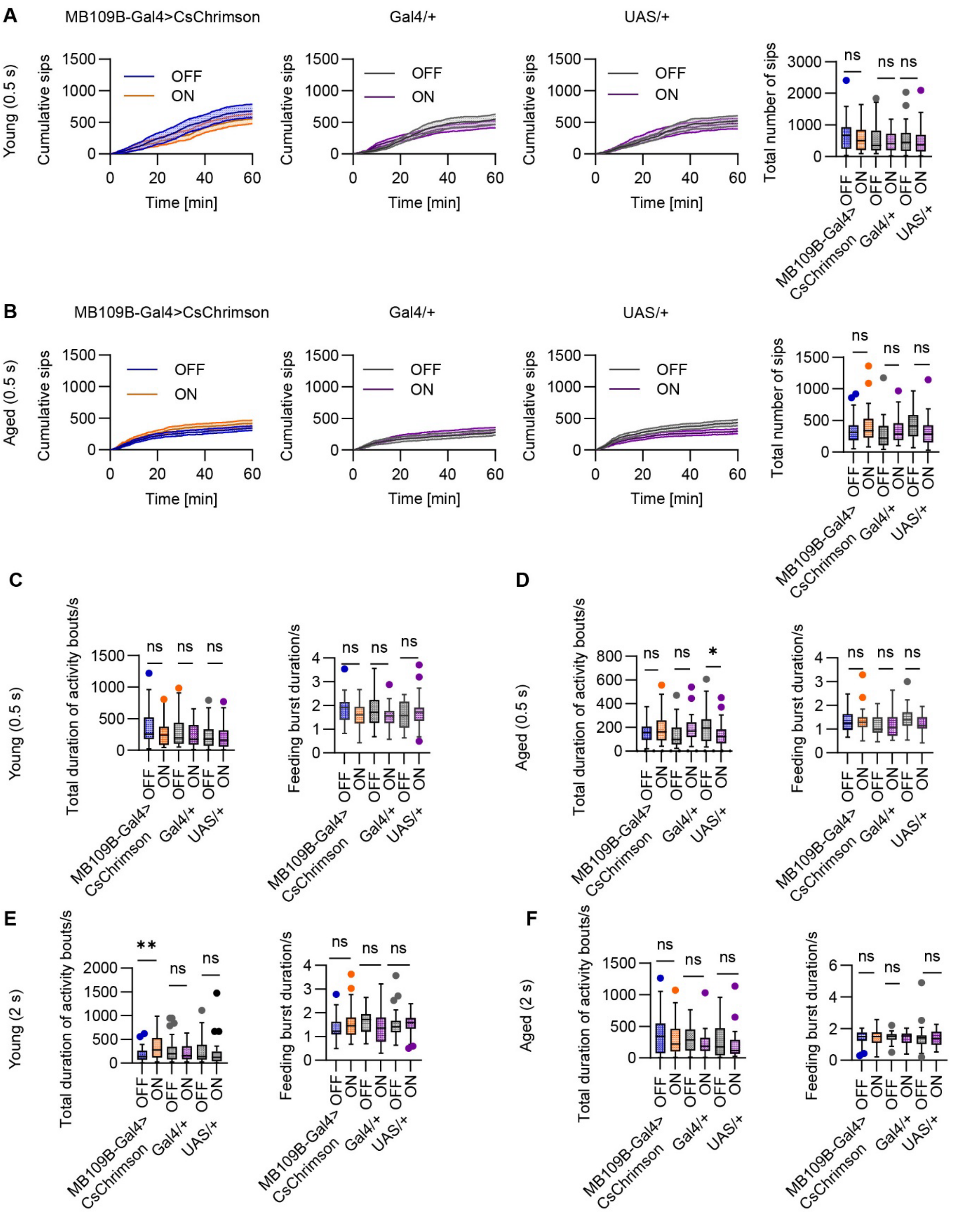


**Figure S3. Feeding behavior and parameters analysis following activation of dopaminergic PAM-β′2a neurons (Related to Figure 4A-C)**

(A–B) Feeding behavior following short optogenetic activation (0.5 s, 5% sucrose) of PAM-β′2a neurons in young (A) and aged (B) flies. Cumulative sip number over the 60 min assay (left) and total sip number at 60 min (right) are shown under light OFF and light ON conditions for MB109B-Gal4>CsChrimson experimental flies (young n = 27, aged n = 32), Gal4/+ controls (young n = 32, aged n = 29), and UAS/+ controls (young n = 31, aged n = 31).

(C–D) Feeding parameters analysis following short optogenetic activation (0.5 s) of PAM-β′2a neurons in young (C) and aged (D) flies. Total duration of activity bouts (left) and feeding burst duration (right) are shown under light OFF and light ON conditions for the indicated genotypes.

(E–F) Feeding parameters analysis following prolonged optogenetic activation (2 s, 2% sucrose) of PAM-β′2a neurons in young (E) and aged (F) flies. Total duration of activity bouts (left) and feeding burst duration (right) are shown under light OFF and light ON conditions for MB109B-Gal4>CsChrimson experimental flies (young $n = 29$, aged $n = 30$), Gal4/+ controls (young $n = 29$, aged $n = 32$), and UAS/+ controls (young $n = 27$, aged $n = 26$).

Light OFF (blue; grey) and light ON (orange; purple). Statistical comparisons were performed using the Wilcoxon matched-pairs signed-rank test. (A) Young (0.5 s): MB109B-Gal4>CsChrimson, $p = 0.4873$; Gal4/+, $p = 0.5515$; UAS/+, $p = 0.5814$. (B) Aged (0.5 s): MB109B-Gal4>CsChrimson, $p = 0.5000$; Gal4/+, $p = 0.3495$; UAS/+, $p = 0.0527$. (C) Young (0.5 s), Total duration of activity bouts: MB109B-Gal4>CsChrimson, $p = 0.3361$; Gal4/+, $p = 0.6511$; UAS/+, $p = 0.5682$. Feeding burst duration: MB109B-Gal4>CsChrimson, $p = 0.0595$; Gal4/+, $p = 0.1870$; UAS/+, $p = 0.1445$. (D) Aged (0.5 s), Total duration of activity bouts: MB109B-Gal4>CsChrimson, $p = 0.3595$; Gal4/+, $p = 0.1494$; UAS/+, $p = 0.0414$. Feeding burst duration: MB109B-Gal4>CsChrimson, $p = 0.9576$; Gal4/+, $p = 0.6201$; UAS/+, $p = 0.1130$. (E) Young (2 s), Total duration of activity bouts: MB109B-Gal4>CsChrimson, $p = 0.0038$; Gal4/+, $p = 0.3927$; UAS/+, $p = 0.2901$. Feeding burst duration: MB109B-Gal4>CsChrimson, $p = 0.0974$; Gal4/+, $p = 0.7664$; UAS/+, $p = 0.1307$. (F) Aged (2 s), Total duration of activity bouts: MB109B-Gal4>CsChrimson, $p = 0.3184$; Gal4/+, $p = 0.0945$; UAS/+, $p = 0.2079$. Feeding burst duration: MB109B-Gal4>CsChrimson, $p = 0.9598$; Gal4/+, $p = 0.2294$; UAS/+, $p = 0.7260$.

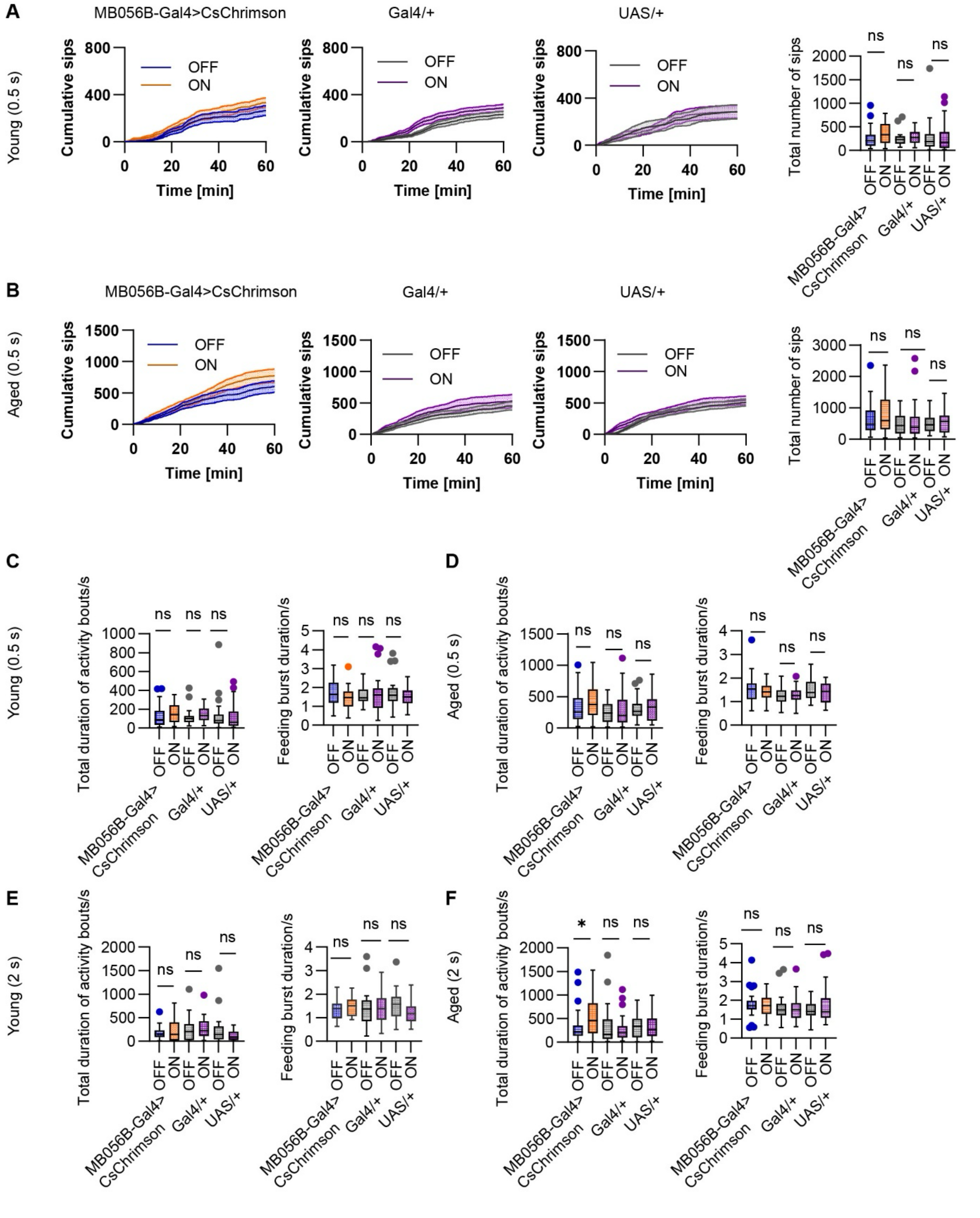


**Figure S4. Feeding behavior and parameters analysis following activation of dopaminergic PAM-β'2m/β'2p neurons (Related to Figure 4D-F)**

(A–B) Feeding behavior following short optogenetic activation (0.5 s, 5% sucrose) of PAM-β'2m/β'2p neurons in young (A) and aged (B) flies. Cumulative sip number over the 60 min assay (left) and total sip number at 60 min (right) are shown under light OFF and light ON conditions for MB056B-Gal4>CsChrimson experimental flies (young n = 28, aged n = 29), Gal4/+ controls (young n = 31, aged n = 29), and UAS/+ controls (young n = 32, aged n = 29).

(C–D) Feeding parameters analysis following short optogenetic activation (0.5 s) of PAM-β'2m/β'2p neurons in young (C) and aged (D) flies. Total duration of activity bouts (left) and feeding burst duration (right) are shown under light OFF and light ON conditions for the indicated genotypes.

(E–F) Feeding parameters analysis following prolonged optogenetic activation (2 s, 2% sucrose) of PAM-β'2m/β'2p neurons in young (E) and aged (F) flies. Total duration of activity bouts (left) and feeding burst duration (right) are shown under light OFF and light ON conditions for MB056B-Gal4>CsChrimson experimental flies (young n = 21, aged n = 28), Gal4/+ controls (young n = 21, aged n = 30), and UAS/+ controls (young n = 21, aged n = 26).

Light OFF (blue; grey) and light ON (orange; purple). Statistical comparisons were performed using the Wilcoxon matched-pairs signed-rank test. (A) Young (0.5 s): MB056B-Gal4>CsChrimson, p = 0.2741; Gal4/+, p = 0.1393; UAS/+, p = 0.9192. (B) Aged (0.5 s): MB056B-Gal4>CsChrimson, p = 0.2404; Gal4/+, p = 0.6089; UAS/+, p = 0.4846. (C) Young (0.5 s), Total duration of activity bouts: MB056B-Gal4>CsChrimson, p = 0.3502; Gal4/+, p = 0.0982; UAS/+, p = 0.8848. Feeding burst duration: MB056B-Gal4>CsChrimson, p = 0.1054; Gal4/+, p = 0.2702; UAS/+, p = 0.9241. (D) Aged (0.5 s), Total duration of activity bouts: MB056B-Gal4>CsChrimson, p = 0.1757; Gal4/+, p = 0.3250; UAS/+, p = 0.3692. Feeding burst duration: MB056B-Gal4>CsChrimson, p = 0.5710; Gal4/+, p = 0.9618; UAS/+, p = 0.0952. (E) Young (2 s), Total duration of activity bouts: MB056B-Gal4>CsChrimson, p = 0.4319; Gal4/+, p = 0.1907; UAS/+, p = 0.1111. Feeding burst duration: MB056B-Gal4>CsChrimson, p = 0.2722; Gal4/+, p = 0.7399; UAS/+, p = 0.1819. (F) Aged (2 s), Total duration of activity bouts: MB056B-Gal4>CsChrimson, p = 0.0340; Gal4/+, p = 0.8236; UAS/+, p = 0.4525. Feeding burst duration: MB056B-Gal4>CsChrimson, p = 0.5865; Gal4/+, p = 0.9556; UAS/+, p = 0.9057.
